# Interpretable detection of novel human viruses from genome sequencing data

**DOI:** 10.1101/2020.01.29.925354

**Authors:** Jakub M. Bartoszewicz, Anja Seidel, Bernhard Y. Renard

**Affiliations:** Bioinformatics (MF1), Department of Methodology and Research Infrastructure, Robert Koch Institute, Berlin, Germany; Department of Mathematics and Computer Science, Free University of Berlin, Berlin, Germany; Data Analytics and Computation Statistics, Hasso Plattner Institute for Digital Engineering, Potsdam, Brandenburg, Germany; Digital Engineering Faculty, University of Postdam, Potsdam, Brandenburg, Germany; Central Research Institute of Ambulatory Health Care, Berlin, Germany

## Abstract

Viruses evolve extremely quickly, so reliable methods for viral host prediction are necessary to safeguard biosecurity and biosafety alike. Novel human-infecting viruses are difficult to detect with standard bioinformatics workflows. Here, we predict whether a virus can infect humans directly from next-generation sequencing reads. We show that deep neural architectures significantly outperform both shallow machine learning and standard, homology-based algorithms, cutting the error rates in half and generalizing to taxonomic units distant from those presented during training. Further, we develop a suite of interpretability tools and show that it can be applied also to other models beyond the host prediction task. We propose a new approach for convolutional filter visualization to disentangle the information content of each nucleotide from its contribution to the final classification decision. Nucleotide-resolution maps of the learned associations between pathogen genomes and the infectious phenotype can be used to detect regions of interest in novel agents, for example the SARS-CoV-2 coronavirus, unknown before it caused a COVID-19 pandemic in 2020. All methods presented here are implemented as easy-to-install packages enabling analysis of NGS datasets without requiring any deep learning skills, but also allowing advanced users to easily train and explain new models for genomics.

## INTRODUCTION

### Background

Within a globally interconnected and densely populated world, pathogens can spread more easily than they ever had before. As the recent outbreaks of Ebola and Zika viruses have shown, the risks posed even by these previously known agents remain unpredictable and their expansion hard to control (1). What is more, it is almost certain that more unknown pathogen species and strains are yet to be discovered, given their constant, extremely fast-paced evolution and unexplored biodiversity, as well as increasing human exposure (2, 3). Some of those novel pathogens may cause epidemics (similar to the SARS and MERS coronavirus outbreaks in 2002 and 2012) or even pandemics (e.g. SARS-CoV-2 and the “swine flu” H1N1/09 strain). Many have more than one host or vector, which makes assessing and predicting the risks even more difficult. For example, Ebola has its natural reservoir most likely in fruit bats (4), but causes deadly epidemics in both humans and chimpanzees. As the state-of-the art approach for the open-view detection of pathogens is genome sequencing (5, 6), it is crucial to develop automated pipelines for characterizing the infectious potential of currently unidentifiable sequences. In practice, clinical samples are dominated by host reads and contaminants, with often less than a hundred reads of the pathogenic virus (7). Metagenomic assembly is challenging, especially in time-critical applications. This creates a need for read-based approaches complementing or substituting assembly where needed.

Screening against potentially dangerous subsequences before their synthesis may also be used as a way of ensuring responsible research in synthetic biology. While potentially useful in some applications, engineering of viral genomes could also pose a biosecurity and biosafety threat. Two controversial studies modified the influenza A/H5N1 (“bird flu”) virus to be airborne transmissible in mammals (8, 9). A possibility of modifying coronaviruses to enhance their virulence triggered calls for a moratorium on this kind of research (10). Synthesis of an infectious horsepox virus closely related to the smallpox-causing *Variola* virus (11) caused a public uproar and calls for intensified discussion on risk control in synthetic biology (12).

### Current tools for host range prediction

Several computational, genome-based methods exist that allow to predict the host-range of a bacteriophage (a bacteria-infecting virus). A selection of composition-based and alignment-based approaches has been presented in an extensive review by Edwards et al. (13). Prediction of eukariotic host tropism (including humans) based on known protein sequences was shown for the influenza A virus (14). Support-vector machines based on word2vec representations were shown to outperform homology searches with BLAST and HMMs in the same task, but lost their advantage when applied to nucleic acid sequences directly (15). Two recent studies employ *k*-mer based, *k*-NN classifiers (16) and deep learning (17) to predict host range for a small set of three well-studied species directly from viral sequences. While those approaches are limited to those particular species and do not scale to viral host-range prediction in general, the Host Taxon Predictor (HTP) (18) uses logistic regression and support vector machines to predict if a novel virus infects bacteria, plants, vertebrates or arthropods. Yet, the authors argue that it is not possible to use HTP in a read-based manner; it requires long sequences of at least 3,000 nucleotides. This is incompatible with modern metagenomic next-generation sequencing (NGS) workflows, where the DNA reads obtained are at least 10-20 times shorter. Another study used gradient boosting machines to predict reservoir hosts and transmission via arthropod vectors for known human-infecting viruses (19).

Zhang et al. (20) designed several classifiers explicitly predicting whether a new virus can potentially infect humans. Their best model, a *k*-NN classifier, uses *k*-mer frequencies as features representing the query sequence and can yield predictions for sequences as short as 500 base pairs (bp). It worked also with 150bp-long reads from real DNA sequencing runs, although in this case the reads originated also from the viruses present in the training set (and were therefore not “novel”).

### Deep Learning for DNA sequences

While DNA sequences mapped to a reference genome may be represented as images (21), a majority of studies uses a distributed orthographic representation, where each nucleotide {*A,C,G,T*} in a sequence is represented by a one-hot encoded vector of length 4. An “unknown” nucleotide (*N*) can be represented as an all-zero vector. Chaos game representation (CGR) and its extension, the frequency matrix CGR (FCGR) are promising alternatives able to encode an arbitrary sequence in an image-like format. FCGR has been used to encode genomic inputs for deep learning approaches, including full bacterial genomes (22) and coding sequences of HIV for the drug resistance prediction task (23). In this study, we use one-hot encoding with *N* s as zeroes, which was previously shown to perform well for raw NGS reads (24) and abstract phenotype labels.

CNNs and LSTMs have been successfully used for a variety of DNA-based prediction tasks. Early works focused mainly on regulation of gene expression in humans (25, 26, 27, 28, 29), which is still an area of active research (30, 31, 32). In the field of pathogen genomics, deep learning models trained directly on DNA sequences were developed to predict host ranges of three multi-host viral species (33) and to predict pathogenic potentials of novel bacteria (24). DeepVirFinder (34) and ViraMiner (35) can detect viral sequences in metagenomic samples, but they cannot predict the host and focus on previously known species. For a broader view on deep learning in genomics we refer to a recent review by Eraslan et al. (36).

Interpretability and explainability of deep learning models for genomics is crucial for their wide-spread adoption, as it is necessary for delivering trustworthy and actionable results. Convolutional filters can be visualized by forward-passing multiple sequences through the network and extracting the most-activating subsequences (25) to create a position weight matrix (PWM) which can be visualized as a sequence logo (37, 38). Direct optimization of input sequences is problematic, as it results in generating a dense matrix even though the input sequences are one-hot encoded (39, 40). This problem can be alleviated with Integrated Gradients (41, 42) or DeepLIFT, which propagates activation differences relative to a selected reference back to the input, reducing the computational overhead of obtaining accurate gradients (43). If the bias terms are zero and a reference of all-zeros is used, the method is analogous to Layer-wise Relevance Propagation (44). DeepLIFT is an additive feature attribution method, and may used to approximate Shapley values if the input features are independent (45). TF-MoDISco (46) uses DeepLIFT to discover consolidated, biologically meaningful DNA motifs (transcription factor binding sites).

### Contributions

In this paper, we first improve the performance of read-based predictions of the viral host (human or non-human) from next-generation sequencing reads. We show that reverse-complement (RC) neural networks (24) significantly outperform both the previous state-of-the-art (20) and the traditional, alignment-based algorithm – BLAST (47, 48), which constitutes a gold standard in homology-based bioinformatics analyses. We show that defining the negative (non-human) class is non-trivial and compare different ways of constructing the training set. Strikingly, a model trained to distinguish between viruses infecting humans and viruses infecting other chordates (a phylum of animals including vertebrates) generalizes well to evolutionarily distant non-human hosts, including even bacteria. This suggests that the host-related signal is strong and the learned decision boundary separates human viruses from other DNA sequences surprisingly well.

Next, we propose a new approach for convolutional filter visualization using partial Shapley values to differentiate between simple nucleotide information content and the contribution of each sequence position to the final classification score. To test the biological plausibility of our models, we generate genome-wide maps of “infectious potential” and nucleotide contributions. We show that those maps can be used to visualize and detect virulence-related regions of interest (e.g. genes) in novel genomes.

As a proof of concept, we analyzed one of the viruses randomly assigned to the test set – the Taï Forest ebolavirus, which has a history of host-switching and can cause a serious disease. To show that the method can also be used for other biological problems, we investigated the networks trained by Bartoszewicz et al. (24) and their predictions on a genome of a pathogenic bacterium *Staphylococcus aureus*. The authors used this particular species to assess the performance of their method on real sequencing data. Finally, we studied the SARS-CoV-2 coronavirus, which emerged in December 2019, causing the COVID-19 pandemic (49).

## MATERIALS AND METHODS

### Data collection and preprocessing

#### VHDB dataset

We accessed the Virus-Host Database (50) on July 31, 2019 and downloaded all the available data. We note that all the reference genomes from NCBI Viral Genomes are present in VHDB, as well as their curated annotations from RefSeq. Additional, manually curated records in VHDB extend on metadata available in NCBI. More non-reference genomes are available, but considering multiple genomes per virus would skew the classifiers’ performance towards the more frequently resequenced ones.

The original dataset contained 14,380 records comprising RefSeq IDs for viral sequences and associated metadata. Some viruses are divided into discontiguous segments, which are represented as separate records in VHDB; in those cases the segments were treated as contigs of a single genome in the further analysis. We removed records with unspecified host information and those confusing the highly pathogenic Variola virus with a similarly named genus of fish. Following Zhang et al. (20), we filtered out viroids and satellites, which are classified as subviral agents and not *bona fide* viruses (51, 52). Note that even though they require helper viruses for replication, this step did not affect ubiquitous adeno-associated viruses and large virophages, which are well established within the viral taxonomy in the families *Parvoviridae* and *Lavidaviridae*, respectively. Human-infecting viruses were extracted by searching for records containing “Homo sapiens” in the “host name” field. Note that VHDB contains information about multiple possible hosts for a given virus where appropriate. Any virus infecting humans was assigned to the positive class, also if other, non-human hosts exist. In total, the dataset contained 9,496 viruses (grouped in 7503 species), including 1,309 human viruses (393 species). We considered both DNA and RNA viruses; RNA sequences were encoded in the DNA alphabet, as in RefSeq.

#### Defining the negative class

While defining a human-infecting class is relatively straightforward, the reference negative class may be conceptualized in a variety of ways. The broadest definition takes all non-human viruses into account, including bacteriophages (bacterial viruses). This is especially important, as most of known bacteriophages are DNA viruses, while many important human (and animal) viruses are RNA viruses. One could expect that the multitude of available bacteriophage genomes dominating the negative class could lower the prediction performance on viruses similar to those infecting humans. This offers an open-view approach covering a wider part of the sequence space, but may lead to misclassification of potentially dangerous mammalian or avian viruses. As they are often involved in clinically relevant host-switching events, a stricter approach must also be considered. In this case, the negative class comprises only viruses infecting Chordata (a group containing vertebrates and closely related taxa). Two intermediate approaches consider all eukaryotic viruses (including plant and fungi viruses), or only animal-infecting viruses. This amounts to four nested host sets: “All” (8,187 non-human viruses, 7110 species), “Eukaryota” (5,114 viruses, 4275 species), “Metazoa” (2,942 viruses, 2351 species) and “Chordata” (2,078 viruses, 1530 species). Auxiliary sets containing only non-eukaryotic viruses (“non-Eukaryota”), non-animal eukaryotic viruses (“non-Metazoa Eukaryota”) etc. can be easily constructed by set subtraction.

For the positive class, we randomly generated a training set containing 80% of the genomes, and validation and test sets with 10% of the genomes each. Importantly, the nested structure was kept also during the training-validation-test split: for example, the species assigned to the smallest test set (“Chordata”) were also present in all the bigger test sets. The same applied to other taxonomic levels, as well as the training and validation sets wherever applicable.

#### Read simulation

We simulated 250bp long Illumina reads following a modification of a previously described protocol (24) and using the Mason read simulator (53). First, we only generated the reads from the genomes of human-infecting viruses. Then, the same steps were applied to each of the four negative class sets. Finally, we also generated a fifth set, “Stratified”, containing an equal number of reads drawn from genomes of the following disjunct host classes: “Chordata” (25%), “non-Chordata Metazoa” (25%), “non-Metazoa Eukaryota” (25%) and “non-Eukaryota” (25%).

In each of the evaluated settings, we used a total of 20 million (80%) reads for training, 2.5 million (10%) reads for validation and 2.5 million (10%) paired reads as the held-out test set. Read number per genome was proportional to genome length, keeping the coverage uniform on average. Viruses with longer genomes were therefore represented by more reads than shorter viruses. On the other hand, their sequence diversity was covered at a similar level. This length-balancing step was previously shown to work well for bacterial genomes of different lengths (24, 54). While the original datasets are heavily imbalanced, we generated the same number of negative and positive data points (reads) regardless of the negative class definition used.

This protocol allowed us to test the impact of defining the negative class, while using the exactly same data as representatives of the positive class. We used three training and validation sets (“All”, “Stratified”, and “Chordata”), representing the fully open-view setting, a setting more balanced with regard to the host taxonomy, and a setting focused on cases most likely to be clinically relevant. In each setting, the validation set matched the composition of the training set. The evaluation was performed using all five test sets to gain a more detailed insight on the effects of negative class definition on the prediction performance.

#### Human blood virome dataset

Similarily to Zhang et al. (20), we used the human blood DNA virome dataset (55) to test the selected classifiers on real data. We obtained 14,242,329 reads of 150bp and searched all of VHDB using blastn (with default parameters) to obtain high-quality reference labels. If a read’s best hit was a human-infecting virus, we assigned it to a positive class; the negative class was assigned if this was not the case. This procedure yielded 14,012,665 “positive” and 229,664 “negative” reads.

#### Virus-level and species-level predictions

In this study, we focus on predicting labels for reads originating from novel viruses. What constitutes a “novel” biological entity is an open question – a novel virus does not necessarily belong to a novel species (56). If a given viral isolate clusters with a known group of isolates, it is considered to be the same virus; if it does not, it may be assigned a distinct name and considered novel (56). This is separate from its putative taxonomical assignment. Assigning a novel virus to a novel or a previously established species is performed pursuing a wider set of criteria, and the criteria for delineating distinct species differ between viral families (51, 52, 56, 57). In most cases, species are perceived as human constructs rather than biological entities and host range often is explicitly one of the defining features (56, 58), rendering reasoning based on cross-species homology searches inherently difficult.

The most prominent example of this problem is the SARS-CoV-2 virus, which is a novel virus within a previously known species (*Severe acute respiratory syndrome–related coronavirus*). Other members of this species include the human-infecting SARS-CoV-1, but also multiple related bat SARSr-CoV viruses (e.g. SARSr-CoV RaTG13 or Bat SARS-like coronavirus WIV1). Importantly, SARS-CoV-2 is not a strain of SARS-CoV-1; those two viruses share a common ancestor (56). This echoes similar problems related to pathogenic potential prediction for novel bacterial pathogens. A novel bacterium may be defined as a novel strain or a novel species (24), and the classifiers must be trained according to the desired definition.

As the 2020 pandemic has shown, different viruses of the same species can differ wildly in their infectious potential and the broader impact on human societies. Therefore, threat assessment must be performed for novel viruses, not only novel taxa; different related viruses are non-redundant. At the same time, redundancy below this level (i.e. multiple instances of the same virus) must be eliminated from the dataset to ensure reliability of the trained classifier. VHDB tackles this problem by collecting and annotating reference genomes – each virus in the database is a separate entity with its own ID in NCBI Taxonomy. This virus-level approach was previously used by Zhang et al. (20). We show that homology-based algorithms underperform in this setting already, suggesting that machine learning is indeed required to accurately predict labels for novel viruses even if other members of the same species are present in the training database.

Nevertheless, a more difficult alternative – predictions for reads of viruses belonging to completely novel species – is a related and potentially equally important task. For bacterial datasets, species novelty can be modelled by selecting a single representative genome per species (24). As the SARS-CoV-2 example shows, this is often not possible for viruses. To assess our approach in this stricter setup, we re-divided the VHDB dataset into training, validation and test sets ensuring that all viruses of a given species were assigned to only one of those subsets. This effectively models a “novel species” scenario while also reflecting within-species phenotype diversity. We recreated the species-wide versions of the “All” and “Chordata” datasets by assigning 80%, 10% and 10% of the species to the training, validation and test datasets, respectively. We resimulated the reads as outlined above and compared the performance of the machine learning and homology-based approaches achieving the highest accuracy in the simpler “novel virus” setting (see Section Prediction performance).

### Training

We used the DeePaC package (24) to investigate RC-CNN and RC-LSTM architectures, which guarantee identical predictions for both forward and reverse-complement orientations of any given nucleotide sequence, and have been previously shown to accurately predict bacterial pathogenicity. Here, we employ an RC-CNN with two convolutional layers with 512 filters of size 15 each, average pooling and 2 fully connected layers with 256 units each. The LSTM used has 384 units (Fig. S1). We use dropout regularization in both cases, together with aggressive input dropout at the rate of 0.2 or 0.25 (tuned for each model). Input dropout may be interpreted as a special case of noise injection, where a fraction of input nucleotides is turned to *N* s. Representations of forward and reverse-complement strands are summed before the fully connected layers. As two mates in a read pair should originate from the same virus, predictions obtained for them can be averaged for a boost in performance. If a contig or genome is available, averaging predictions for constituting reads yields a prediction for the whole sequence. We used Tesla P100 and Tesla V100 GPUs for training and an RTX 2080 Ti for visualizations.

We wanted the networks to yield accurate predictions for both 250bp (our data, modelling a sequencing run of an Illumina MiSeq device) and 150bp long reads (as in the Human Blood Virome dataset). As shorter reads are padded with zeros, we expected the CNNs trained using average pooling to misclassify many of them. Therefore, we prepared a modified version of the datasets, in which the last 100bp of each read were turned to zeros, mocking a shorter sequencing run while preserving the error model. Then, we retrained the CNN which had performed best on the original dataset. Since in principle, the Human Blood Virome dataset should not contain viruses infecting non-human Chordata, a “Chordata”-trained classifier was not used in this setting.

### Benchmarking

We compare our networks to the the *k*-NN classifier proposed by Zhang et al. (20), the only other approach explicitly tested on raw NGS reads and detecting human viruses in a fully open view setting (not focusing on a limited number of species). We use the real sequencing data that they used (55) for an unbiased comparison.

We trained the classifier on the “All” dataset as described by the authors, i.e. using non-overlapping, 500bp-long contigs generated from the training genomes (retraining on simulated reads is computationally prohibitive). We also tested the performance of using BLAST to search against an indexed database of labeled genomes. We constructed the database from the “All” training set and used discontiguous megablast to achieve high inter-species sensitivity. For NGS mappers (BWA-MEM (59) and Bowtie2 (60)), the indices were constructed analogously. Kraken (61) was previously shown to perform worse than both BLAST and machine learning when faced with read-based pathogenic potential prediction for novel bacterial species (54). Its major advantage – assigning reads to lowest common ancestor (LCA) nodes in ambiguous cases – turns into a problem in the infectivity prediction task, as transferring labels to LCAs is often impossible (54). Therefore, we focus on alignment-based approaches as the most accurate alternative to machine learning in this context.

Note that both alignment and *k*-NN can yield conflicting predictions for the individual mates in a read pair. What is more, BLAST and the mappers yield no prediction at all if no match is found. Therefore, similarly to Bartoszewicz et al. (24), we used the *accept anything* operator to integrate binary predictions for read pairs and genomes. At least one match is needed to predict a label, and conflicting predictions are treated as if no match was found at all. Missing predictions lower both true positive and true negative rates.

### Filter visualization

#### Substring extraction

In order to visualize the learned convolutional filters, we downsample a matching test set to 125,000 reads and pass it through the network. This is modelled after the method presented by Alipanahi et al. (25). For each filter and each input sequence, the authors extracted a subsequence leading to the highest activation, and created sequence logos from the obtained sequence sets (“max-activation”). We used the DeepSHAP implementation (45) of DeepLIFT (43) to extract score-weighted subsequences with the highest contribution score (“max-contrib”) or all score-weighted subsequences with non-zero contributions (“all-contrib”). Computing the latter was costly and did not yield better quality logos.

We use an all-zero reference. As reads from real sequencing runs are usually not equally long, shorter reads must be padded with *N* s; the “unknown” nucleotide is also called whenever there is not enough evidence to assign any other to the raw sequencing signal. Therefore, *N* s are “null” nucleotides and are a natural candidate for the reference input. We do not consider alternative solutions based on GC content or dinucleotide shuffling, as the input reads originate from multiple different species, and the sequence composition may itself be a strong marker of both virus and host taxonomy (13). We also avoid weight-normalization suggested for zero-references (43), as it implicitly models the expected GC content of all possible input sequences, and assumes no *N* s present in the data. Finally, we calculate average filter contributions to obtain a crude ranking of feature importance with regard to both the positive and negative class.

#### Partial Shapley values

Building sequence logos involves calculating information content (IC) of each nucleotide at each position in a prospective DNA motif. This can be then interpreted as measure of evolutionary sequence conservation. However, high IC does not necessarily imply that a given nucleotide is relevant in terms of its contribution to the classifier’s output. Some sub-motifs may be present in the sequences used to build the logo, even if they do not contribute to the final prediction (or even a given filter’s activation).

To test this hypothesis, we introduce partial Shapley values. Intuitively speaking, we capture the contributions of a nucleotide to the network’s output, but only in the context of a given intermediate neuron of the convolutional layer. More precisely, for any given feature *x*_*i*_, intermediate neuron *y*_*j*_ and the output neuron *z*, we aim to measure how *x*_*i*_ contributes to *z* while regarding only the fraction of the total contribution of *x*_*i*_ that influences how *y*_*j*_ contributes to *z*. Although similarly named concepts were mentioned before as intermediate computation steps in a different context (62, 63), we define and use partial Shapley values to visualize contribution flow through convolutional filters. This differs from recently introduced contribution weight matrices (32), where feature attributions are used as a representation of an identified transcription factor binding site irreducible to a given intermediate neuron.

Using the formalism of DeepLIFT’s multipliers (43) and their reinterpretation in SHAP (45), we backpropagate the activation differences only along the paths “passing through” *y*_*j*_ In Eq. **1**, we define partial multipliers 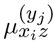 and express them in terms of Shapley values *ϕ* and activation differences w.r.t. the expected activation values (reference activation). Calculating partial multipliers is equivalent to zeroing out the multipliers 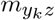 for all *k* ≠ *j* before backpropagating 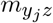 further. 

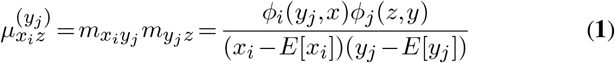

We define partial Shapley values 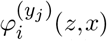 analogously to how Shapley values can be approximated by a product of multipliers and input differences w.r.t. the reference (Eq. **2**): 

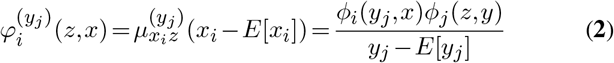

From the chain rule for multipliers (43), it follows that standard multipliers are a sum over all partial multipliers for a given layer *y*. Therefore, Shapley values as approximated by DeepLIFT are a sum of partial Shapley values for the layer *y* (Eq. **3**). 

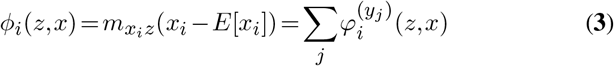

Once we calculate the contributions of convolutional filters for the first layer, 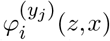 for the first convolutional layer of a network with one-hot encoded inputs and an all-zero reference can be efficiently calculated using weight matrices and filter activation differences (Eq. **4**-**5**). First, in this case we do not traverse any non-linearities and can directly use the linear rule (43) to calculate the contributions of *x*_*i*_ to *y*_*j*_ as a product of the weight *w*_*i*_ and the input *x*_*i*_. Second, the input values may only be 0 or 1. 

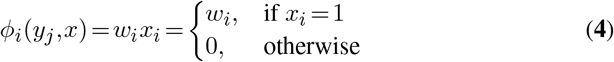

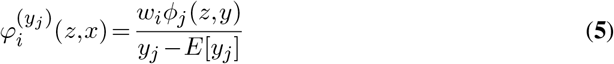

Resulting partial contributions can be visualized along the IC of each nucleotide of a convolutional kernel. To this end, we design extended sequence logos, where each nucleotide is colored according to its contribution. Positive contributions are shown in red, negative contributions are blue, and near-zero contributions are gray. Therefore, no information is lost compared to standard sequence logos, but the relevance of individual nucleotides and the filter as a whole can be easily seen. Color saturation is limited by the reciprocal of a user-defined gain parameter, here set to *nm*, where *n* equals the number of input features *x*_*i*_ (sequence length) and *m* equals the number of convolutional filters *y*_*j*_ in a given layer.

### Genome-wide phenotype analysis

We create genome-wide phenotype analysis (GWPA) plots to analyse which parts of a viral genome are associated with the infectious phenotype. We scramble the genome into overlapping, 250bp long subsequences (pseudo-reads) without adding any sequencing noise. For the highest resolution, we use a stride of one nucleotide. For *S. aureus*, we used a stride of 125bp. We predict the infectious potential of each pseudo-read and average the obtained values at each position of the genome. Analogously, we calculate average contributions of each nucleotide to the final prediction of the convolutional network. Finally, we normalize raw infectious potentials into the [−0.5,0.5] interval for a more intuitive graphical representation. We visualize the resulting nucleotide-resolution maps with IGV (64). For protein structures, we average the scores codon-wise to obtain contribution scores per amino acid and visualize them with PyMOL (65).

For well-annotated genomes, we compile a ranking of genes (or other genomic features) sorted by the average infectious potential within a given region. In addition to that, we scan the genome with the learned filters of the first convolutional layer to find genes enriched in subsequences yielding non-zero filter activations. We use Gene Ontology to connect the identified genes of interest with their molecular functions and biological processes they are engaged in.

## RESULTS

### Negative class definition

Choosing which viruses should constitute the negative class is application dependent and influences the performance of the trained models. Table S1 summarizes the prediction accuracy for different combinations of the training and test set composition. The models trained only on human and Chordata-infecting viruses maintain similar, or even better performance when evaluated on viruses infecting a much broader host range, including bacteria. This suggests that the learned decision boundary separates human viruses from all the others surprisingly well. We hypothesize that the human host signal must be relatively strong and contained within the Chordata host signal. Dropout rate of 0.2 resulted in the highest validation accuracy for CNN_Str-150_ and LSTM_Str_. A rate of 0.25 was selected for the other models.

Adding more diversity to the negative class may still boost performance on more diverse test sets, as in the case of CNN trained on the “All” dataset (CNN_All_). This model performs a bit worse on viruses infecting hosts related to humans, but achieves higher accuracy than the “Chordata”-trained models and the best recall overall. Rebalancing the negative class using the “Stratified” dataset helps to achieve higher performance on animal viruses while maintaing high overall accuracy. The LSTMs are outperformed by the CNNs, but they can be used for shorter reads without retraining (see Sections Training and Prediction performance).

### Prediction performance

We selected LSTM_All_ and CNN_All_ for further evaluation. We used a single consumer-grade RTX 2080 Ti GPU to measure inference speed. The CNN classifies 5000 reads/s and the LSTM 1855 reads/s. Analyzing ten million reads takes only 33 minutes using the faster model; linear speed-ups are possible if more GPUs are available. Therefore, the trained models achieve high-throughputs necessary to analyze NGS datasets. Table 1 presents the results of a benchmark using the “All” test set. Low performance of the *k*-NN classifier (20) is caused by frequent conflicting predictions for each read in a read pair. In a single-read setting it achieves 75.5% accuracy, while our best model achieves 87.8% (Table S2). Although BLAST achieves high precision, it yields no predictions for over 10% of the samples. CNN_All_ is the most sensitive and accurate. As expected, standard mapping approaches (BWA-MEM and Bowtie2) struggle with analysing novel pathogens – they are the most precise but the least sensitive. Our approach outperforms them by 15-30%.

**Table 1.** Classification performance in the fully open-view setting (all virus hosts), read pairs. Acc. – accuracy, Prec. – precision, Rec. – recall, Spec. – specificity. Bowtie2, BWA-MEM and BLAST yield no predictions for over 35%, 19% and 10% of the samples, respectively. Best performance in bold.

|  | ACC. | PREC. | REC. | SPEC. |
| --- | --- | --- | --- | --- |
| CNN<sub>ALL</sub> (OURS) | **89.9** | 93.9 | **85.4** | **94.4** |
| LSTM<sub>ALL</sub> (OURS) | 86.4 | 89.0 | 83.0 | 89.8 |
| *k*-NN | 57.1 | 57.8 | 52.1 | 62.0 |
| BOWTIE2 | 58.6 | **99.2** | 59.2 | 58.0 |
| BWA-MEM | 72.8 | 98.9 | 73.9 | 71.8 |
| BLAST | 80.6 | 98.4 | 79.1 | 82.2 |

Although we focus on the extreme case of read-based predictions, our method can also be used on assembled contigs and full genomes if they are available, as well as on read sets from pure, single-virus samples. We note that assembly itself does not yield any labels and a follow-up analysis (via alignment, machine learning or other approaches) is required to correctly classify metagenomic contigs in any case. We ran predictions on contigs without any size filtering with both *k*-NN and BLAST (Table 2). We present performance measures for both individual contigs and whole genome predictions based on contig-wise majority vote. We compare them to BLAST with read-wise majority vote (54) and to read-wise average predictions of our networks, analogous to presented previously for bacteria (24). Our method outperforms BLAST by 1.2% and *k*-NN by 8.9%, even though they have access to the full biological context (full sequences of all contigs in a genome), while we simply average outputs for short reads originating from the contigs.

**Table 2.** Classification performance, all hosts. Whole available genomes. Negative class is the majority class. BAcc. – balanced accuracy, Rec. – recall, Spec. – specificity. BLAST (reads) and our networks use read-wise majority vote or output averaging to aggregate predictions over all reads from a genome. *k*-NN (genome) and BLAST (genome) use contig-wise majority vote. *k*-NN (contigs) and BLAST (contigs) represent performance on individual contigs treated as separate entities. *k*-NN (reads) was not used, as high conflicting prediction rates made read-wise aggregation impracticable.

|  | BACC. | AUPR | REC. | SPEC. |
| --- | --- | --- | --- | --- |
| CNN<sub>ALL</sub> (OURS) | **91.7** | **91.2** | 89.3 | 94.2 |
| LSTM<sub>ALL</sub> (OURS) | 86.3 | 85.8 | **96.2** | 76.4 |
| BLAST (READS) | 90.3 | N/A | 85.5 | **95.1** |
| *k*-NN (GENOME) | 82.8 | N/A | 93.9 | 71.6 |
| BLAST (GENOME) | 90.5 | N/A | 86.3 | 94.6 |
| *k*-NN (CONTIGS) | 83.0 | N/A | 94.3 | 71.6 |
| BLAST (CONTIGS) | 88.4 | N/A | 87.1 | 89.7 |

We benchmarked our models against the human blood virome dataset used by Zhang et al. (20). Our models outperform their *k*-NN classifier. As the positive class massively outnumbers the negative class, all models achieve over 99% precision. CNN_All-150_ performs best (Table 3). However, the positive class is dominated by viruses which are not necessarily novel. The CNN was more accurate on training data, so we expected it to detect those viruses easily.

**Table 3.** Classification performance on the human blood virome dataset. Positive class is the majority class. BAcc. – balanced accuracy, Rec. – recall, Spec. – specificity.

|  | BACC. | AUPR | REC. | SPEC. |
| --- | --- | --- | --- | --- |
| CNN<sub>ALL-150</sub> (OURS) | **96.8** | **>99.9** | **97.3** | **96.2** |
| LSTM<sub>ALL</sub> (OURS) | 91.8 | **>99.9** | 88.2 | 95.5 |
| *k*-NN | 83.1 | 99.5 | 80.9 | 85.4 |

Finally, we repeated the analysis in the “novel species” scenario. Classifying novel viral species when restricted to Chordata-infecting viruses is too challenging for practical purposes (Table S3). Read-wise predictions are not much better than random guesses for both BLAST and CNNs. Low precision of BLAST shows that it often recovers wrong labels even when it does find a match – sequence similarity is not a reliable predictor of the infectious potential in this setting. Even if a whole genome is available, overall accuracy is low. This looks very differently in the fully-open view scenario (Table 4). The CNN trained on the species-wise division of the “All” dataset (CNNSP-All) outperforms BLAST by a wide margin on both reads and genomes. Strikingly, CNN_SP-All_ predictions based on a single read pair achieve higher accuracy than BLAST predictions using whole genomes, mainly due to their significantly higher recall. What is more, pooling predictions from all the reads originating from a given genome does not improve overall CNN_SP-All_ accuracy any further. As CNN_SP-All_ does not reliably outperform its Chordata-trained analog on the “Chordata” dataset (CNN_SP-Cho_, Table S3), we suspect that its relatively high accuracy on the “All” dataset is caused by its high sensitivity while maintaining good specificity on non-Chordata viruses.

**Table 4.** Classification performance, novel species. Top: paired reads (see Table 1). BLAST yields predictions for only 64.3% of the pairs. Bottom: whole available genomes or contigs – negative class is the majority class (see Table 2). BAcc. – balanced accuracy (equal to accuracy for the balanced paired-read dataset), Rec. – recall, Spec. – specificity. BLAST (reads) and our networks use read-wise majority vote or output averaging to aggregate predictions over all reads from a genome. BLAST (genome) uses contig-wise majority vote. BLAST (contigs) represents performance on individual contigs treated as separate entities. Note that low precision is heavily affected by class imbalance.

|  | BACC. | PREC. | REC. | SPEC. |
| --- | --- | --- | --- | --- |
| CNN<sub>SP-ALL</sub> (OURS) | **77.3** | 86.2 | **65.0** | **89.6** |
| BLAST | 47.1 | **94.1** | 17.8 | 76.4 |
| CNN<sub>SP-ALL</sub> (OURS) | **76.8** | 34.2 | **69.8** | 83.8 |
| BLAST (READS) | 61.8 | **46.8** | 30.2 | **93.5** |
| BLAST (GENOME) | 64.0 | 44.9 | 36.5 | 91.5 |
| BLAST (CONTIGS) | 57.9 | 37.9 | 33.6 | 82.1 |

### Filter visualization

Over 84% of all contributing first-layer filters in CNN_All_ have positive average contribution scores. We comment more on this fact in Section Nucleotide contribution logos. For CNN_All_, the average information content of our motifs is strongly correlated nucleotide-wise with IC of DeepBind-like logos (Spearman’s *ρ>* 0.95, *p*< 10^−15^ for all contributing filter pairs except one). The difference in average IC is negligible (0.04 bit higher for “max-contrib”, Wilcoxon test, *p*< 10^−15^). Therefore, our contribution logos represent analogous “motifs”, while extracting additional, nucleotide-level interpretations. For exactly one filter, “max-contrib” and “max-activation” scores are not correlated. A deeper analysis reveals that this particular filter is activated by stretches of 0s (*N* s) – it is the only filter with a positive bias, and almost all of its weights are negative (with one near-zero positive). Therefore, an overwhelming majority of its maximum activations are in fact padding artifacts. On the other hand, regions of unambiguous nucleotide sequences result in high positive contributions, since they correspond to a lack of filter activation, where an activation is present for the all-*N* reference. In fact, for over 99.9% of the reads, positive contributions occur at every single position. We suspect that the filter works as an “ambiguity detector”. Since *N* s are modelled as all-zero vectors in the one-hot encoding scheme used here, the network represents “meaningful” (i.e. unambiguous) regions of the input as a missing activation of the filter. This is supported by the fact that the filter lacks any further preference for the specific non-zero nucleotide type. Since sequence logos presented here ignore ambiguous (i.e. noninformative) nucleotides, their ICs for this filter are near-zero, preventing meaningful visualization. On the other hand, this ambiguity seems to play a role in the final classification decision, as contribution distributions are well-separated for both classes (Fig. S2). We speculate that this could be caused by lower quality of the non-pathogen reference genomes, but understanding how exactly this information is used would require further investigation, including feature interactions at all layers of the network. Importantly, only the contribution analysis reveals the relevance of the filter beyond simple activation and nucleotide overrepresentation. The choice of the reference input is crucial.

In the Fig. 1 we present example filters, visualized as “max-contrib” sequence logos based on mean partial Shapley values for each nucelotide at each position. All nucleotides of the filters with the second-highest (Fig. 1a) and the lowest (Fig. 1b) score have relatively strong contributions in accordance with the filters’ own contributions. However, we observe that some nucleotides consistently appear in the activating subsequences, but the sign of their contributions is opposite to the filter’s (low-IC nucleotides of a different color, Fig. 1c). Those “counter-contributions” may arise if a nucleotide with a negative weight forms a frequent motif with others with positive weights strong enough to activate the filter. We comment on this fact in the Section Nucleotide contribution logos. Some filters seem to learn gapped motifs resembling a codon structure (Fig. 1c). We extracted this filter from the original DeePaC network predicting bacterial pathogenicity (24) where the counter-contributions are common, but we find similar filters in our networks as well (Fig. S3). We scanned a genome of *S. aureus* subsp. *aureus* 21200 (RefSeq assembly accession: GCF_000221825.1) with this filter and discovered that the learned motif is indeed significantly enriched in coding sequences (Fisher exact test with Benjamini-Hochberg correction, *q* < 10^−15^). It is also enriched in a number of specific genes. The one with the most hits (sraP, *q* < 10^−15^) is a serine-rich adhesin involved in the pathogenesis of infective endocarditis and mediating binding to human platelets (66). The filter seems to detect serine and glycine repeats in this particular gene (Fig. S5), but a broader, cross-species, multi-gene analysis would be required to fully understand its activation patterns. An analogous analysis revealed that the second-highest contributing filter (Fig. 1a) is overall enriched in coding sequences in both Taï Forest ebolavirus (*q* < 10^−15^, RefSeq accession: NC_014372) and SARS-CoV-2 coronavirus (*q* = 5.6 *×* 10^−5^, RefSeq accession: NC_045512.2). The top hits are the nucleocapsid (N) protein gene of SARS-CoV-2 and the VP35 ebolavirus gene encoding a polymerase cofactor suppressing innate immune signaling (*q* < 10^−15^).

**Figure 1.**
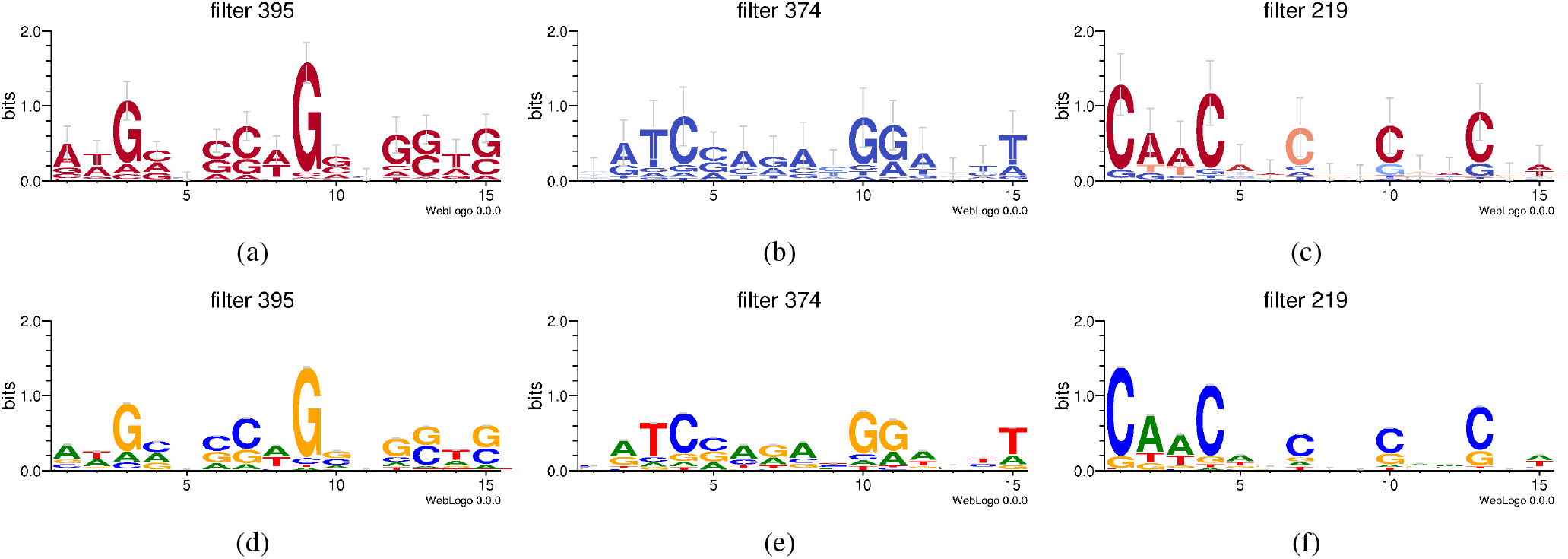
Nucleotide contribution logos of example filters. 1a: Second-highest mean contribution score (CNN_All_). Error bars correspond to Bayesian 95% confidence intervals. 1b: Lowest mean contribution score (CNN_All_). 1c: Gaps resembling a codon structure, extracted from Bartoszewicz et al. (24). Consensus sequence: CAWCNNCNNCNNCNN. 1d-1f: Analogous logos created with the DeepBind-like “max-activation” approach. Our “max-contrib” logos visualize contributions of individual nucleotides, including counter-contributions.

### Genome-wide phenotype analysis

We created a GWPA plot for the Taï Forest ebolavirus genome. Most genes (6 out of 7) can be detected with visual inspection by finding peaks of elevated infectious potential score predicted by at least one of the models (Fig. 2a). Intergenic regions are characterized by lower mean scores. Noticeably, most nucleotide contributions are positive, and low non-negative contributions coincide with regions of negative predictions. Taken together with the surprisingly good generalization of Chordata-trained classifiers and a dominance of positive filters discussed above, this suggests that our networks work as positive class detectors, treating all other sequences as “negative” by default. Indeed, the reference sequence of all *N* s is predicted to be “non-pathogenic” with a score of 0.

**Figure 2.**
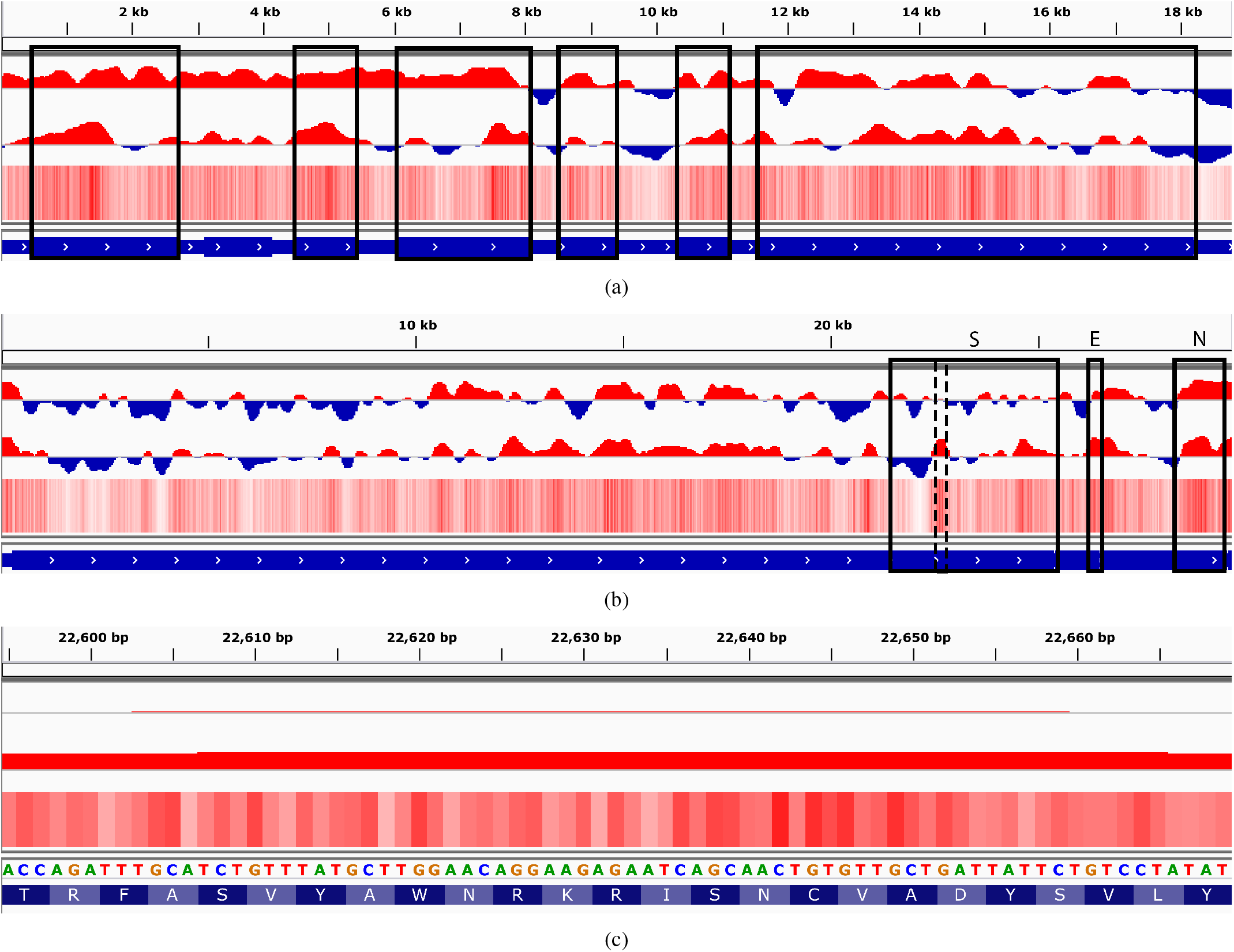
Taï Forest ebolavirus and SARS-CoV-2 coronavirus genomes. Top: score predicted by LSTM_All_. Middle: score predicted by CNN_All_. Heatmap: nucleotide contributions of CNN_All_. Bottom, in blue: reference sequence. 2a: Taï Forest ebolavirus. Genes that can be detected by at least one model are highlighted in black. 2b: Whole genome and sequences encoding the spike protein (S), envelope protein (E) and nucleocapsid protein (N). 2c: Spike protein gene, a small peak (positions 22,595-22,669, dashed line in Fig. 2b) within the receptor-binding domain (predicted by CD-search, positions 22,517-23,185). Binding to the receptor is crucial for entry to the host cell. Local host adaptation could help switch hosts between the animal reservoir and humans.

We ran a similar analysis of *S. aureus* using the built-in DeePaC models (24) and our interpretation workflow. While a viral genome contains usually only a handful of genes, by compiling a ranking of 870 annotated genes of the analyzed *S. aureus* strain we could test if the high-ranking regions are indeed associated with pathogenicity (Table S4). Indeed, out of three top-ranking genes with known biological names and Gene Ontology terms, sarR and sspB are directly engaged in virulence, while hupB regulates expression of virulence-involved genes in many pathogens (67). In contrast to the viral models, both negative and positive contributions are present (Fig. S6), and the model’s output for the all-*N* reference is slightly above the decision threshold (0.58). Even though the network architecture of the viral and the bacterial model are the same, the latter learns a “two-sided” view of the data. We assume this must be a feature of the dataset itself.

Fig. 2b presents a GWPA plot for the whole genome of the SARS-CoV-2 coronavirus, successfully predicted to infect humans, even though the data was collected at least 5 months before its emergence. Interestingly, its mean infectious potential (0.57 as scored by CNN_All_) is relatively close to the decision threshold, while its closest known relative, a bat-infecting SARSr-CoV RaTG13, is actually falsely classified as a human virus with a slightly lower mean infectious potential (0.55). What is more, the gene encoding the spike protein, which plays a significant role in host entry (68), has a mean score slightly above the threshold for SARS-CoV-2 (0.52) and below the threshold for RaTG13 (0.49). As shown in the GWPA plots of both viruses (Fig. 2b and Fig. S4), regions that the network has learned to associate with the infectious phenotype are distributed non-uniformly and tend to cluster together. This suggests that low-confidence mean prediction for those viruses is not a result of random guessing, but genuine ambiguity present in the data – and the misclassification of RaTG13 could be indicative of a general zoonotic potential of SARS-related coronaviruses. In the Fig. 2b, we highlighted the score peaks aligning the spike protein gene (S), as well as the E and N genes, which were scored the highest (apart from an unconfirmed ORF10 of just 38aa downstream of N) by the CNN and the LSTM, respectively. Correlation between the CNN and LSTM outputs is significant, but species-dependent and moderate (0.28 for Ebola, 0.48 for SARS-CoV-2), which suggests they capture complementary signals.

Fig. 2c shows the nucleotide-level contributions in a small peak within the receptor-binding domain (RBD) of the S protein, crucial for recognizing the host cell. The domain location was predicted with CD-search (69) using the default parameters. The maximum score of this peak is noticeably higher for SARS-CoV-2 (0.87) than for its analog in RaTG13 (0.67). Fig. 3 presents the RBD in the structural context of the whole S protein (PDB ID: 6VSB, (70)), as well as in complex with a SARS-neutralizing antibody CR3022 (PDB ID: 6W41, (71)). The high score peak roughly corresponds to one of the regions associated with reduced expression of the RBD (72), located in the core-RBD subdomain. It covers over 71% of the CR3022 epitope, as well as the neighbouring site of the N343 glycan. The latter is present in the epitope of another core-RBD targeting antibody, S309 (73). All the per-residue average contributions in the region are positive (Fig. S7), even in the regions of lower pathogenicity score, in accordance with the results presented in Fig. 2c.

**Figure 3.**
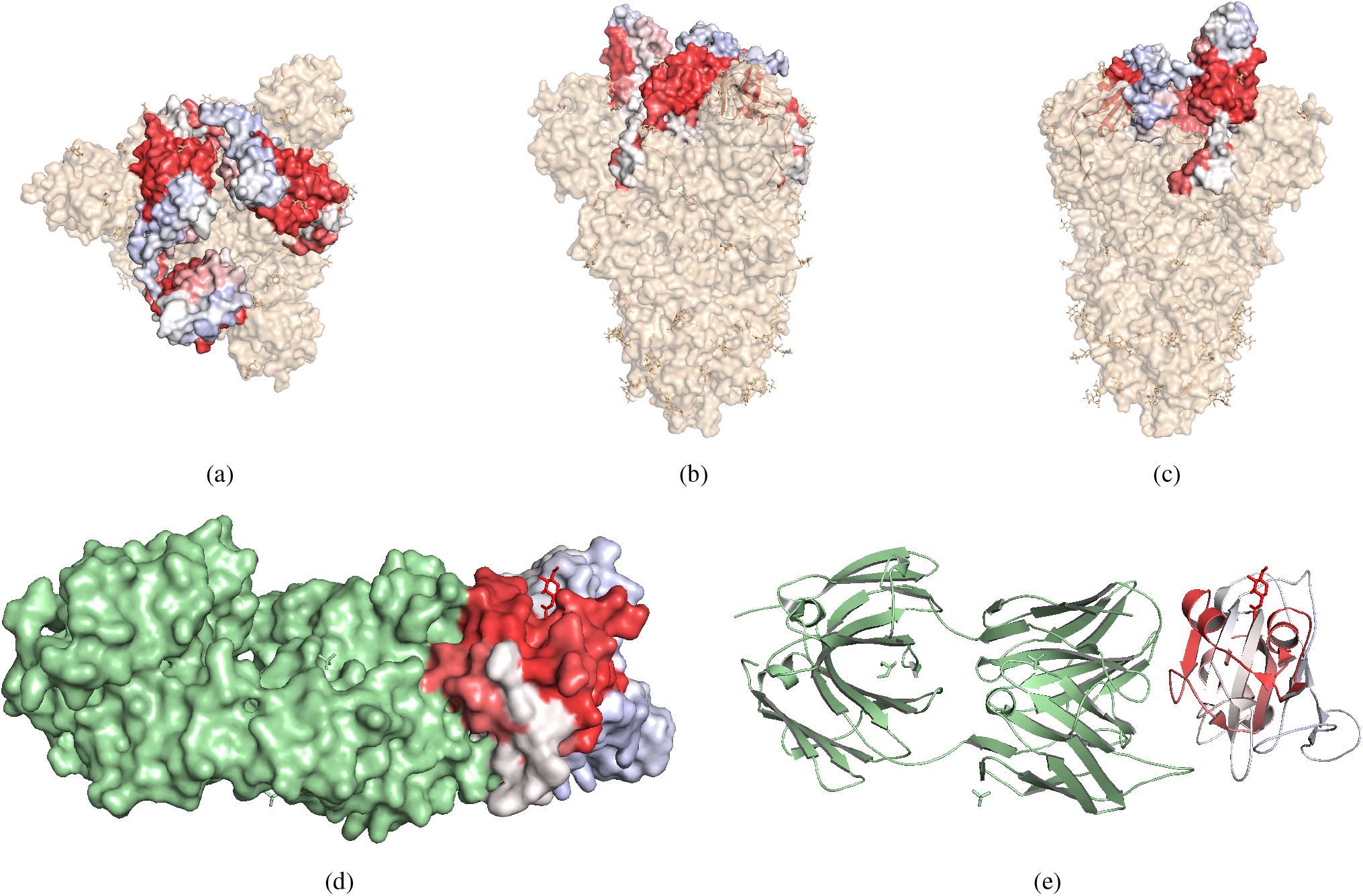
Predicted infectious potentials plotted over the SARS-CoV-2 spike glycoprotein receptor-binding domain. 3a-3c: Top and side view of the spike protein. Three receptor-binding domains (RBDs) are colored in blue, white and red according to the predicted infectious potential of the corresponding genomic sequence. One of the domains is in the “up” conformation. Red regions corresponding to the peak in Fig. 2c are located in the core-RBD subdomain. 3d: RBD in complex with a SARS-neutralizing antibody CR3022 (green). The red region covers over 71% of the CR3022 epitope, but spans also to the neighbouring fragments, including the site of the N343 glycan (carbohydrate in red stick representation). This is a part of the epitope of another neutralizing antibody, S309. 3e: Cartoon representation of Fig. 3d. The red region is centered on two exposed α-helices surrounding the core β-sheet (lower score, white).

## DISCUSSION

### Accurate predictions from short DNA reads

Compared to the previous state-of-the-art in viral host prediction directly from next-generation sequencing reads (20), our models drastically reduce the error rates. This holds also for novel viruses not present in the training set. Generalization of virus-level Chordata models to other host groups is a sign of a strong, “human” signal. We suspect our classifiers detect the positive class treating all other regions of the sequence space as “negative” by default, exhibiting traits of a one-class classifier even without being explicitly trained to do so. We find further support for this hypothesis: the networks learn many more “positive” than “negative” filters and regions of near-zero nucleotide contributions (including the null reference sample) result in negative predictions. As this effect does not occur for bacteria, we expect it do be task- and data-dependent. While we ignore the simulated quality information here, investigating the role of sequencing noise will be an interesting follow-up study. Although the data setup is crucial in general, the modelling step is also important, as shown by our comparison to the baseline *k*-NN model. The RC-nets are relatively simple, but they are invariant to reverse-complementarity and perform better than random forests, naïve Bayes classifiers and standard NN architectures in another NGS task (24).

In the paired read scenario, the previously described *k*-NN approach fails, and standard, alignment-based homology testing algorithms cannot find any matches in more than 10% of the cases, resulting in relatively low accuracy. On a real human virome sample, where a main source of negative class reads is most likely contamination (55), our method filters out non-human viruses with high specificity. In this scenario, the BLAST-derived ground-truth labels were mined using the complete database (as opposed to just a training set). In all cases, our results are only as good as the training data used; high quality labels and sequences are needed to develop trustworthy models. Ideally, sources of error should be investigated with an in-depth analysis of a model’s performance on multiple genomes covering a wide selection of taxonomic units. This is especially important as the method assumes no mechanistic link between an input sequence and the phenotype of interest, and the input sequence constitutes only a small fraction of the target genome without a wider biological context. Still, it is possible to predict a label even from those small, local fragments. A similar effect was also observed for image classification with CNNs (74). Virulence arises as a complex interplay between the host and the virus, so the predictions reflect only an estimated potential of the infectious phenotype. This mirrors the caveats of bacterial pathogenic potential prediction (24), including the considerations of balancing computational cost, reliability of error estimates, size and composition of the reference database. Even though deep learning outperforms the standard homology-based methods, it is still an open question whether it captures “functional” signals, or just a more flexible sequence similarity function. By the very nature of machine learning and sequence comparison in general, we expect similar viruses to yield similar predictions; in principle this could be used to asses a risk of a host-switching event. The interpretability suite presented here aims at shedding some light on this question, but more research is needed.

### Dual-use research and biosecurity

While we focused on the NGS-based prediction scenario, our models could in principle be used to screen DNA synthesis orders for potentially dangerous sequences the context of cyberbiosecurity in synthetic biology. Since standard, homology-based approaches like BLAST are not enough to guarantee accurate screening at a reasonable cost (75, 76, 77), machine learning methods are a promising solution. This has been suggested before for the bacterial DeePaC models (24), and is applicable to the viral networks presented here as well.

However, this line of research can raise questions about possible dual-use. O’Brien and Nelson (78) suggested that while the intended purpose of pathogenicity potential prediction is to mitigate biosecurity threats, it could actually enable designing new pathogens to cause maximal harm. The importance of this concern is difficult to overstate and it must be addressed. If an ML-guided, genome-wide phenotype optimization tool existed, it would indeed be a classical dual-use technology not unlike more established computer-aided design approaches for synthetic biology – potentially dangerous, but offering tremendous benefits (e.g. in agriculture, medicine or manufacturing) as well. However, the models presented here do not allow biologically sensible optimization of target sequences. For example, we find meaningless, low-complexity sequences of mononucleotide repeats corresponding to global maxima (infectious potential of 1.0). These artifacts highlight the fact that only some generally undefined regions of the theoretically possible sequence space are biologically relevant. What is more, we operate on short sequences constituting minuscule fractions of the whole genome with all its complexity. Although successful deep learning approaches for both protein (79, 80, 81) and regulatory sequence design (82, 83, 84, 85) do exist, moving from read-based classification to genome-wide phenotype optimization would require considerable research effort, if possible at all. This would entail capturing a wealth of biological contexts well beyond the capabilities of even the best classification models currently available.

### Nucleotide contribution logos

Visualizing convolutional filters may help to identify more complex filter structures and disentangle the contributions of individual nucleotides from their “conservation” in contributing sequences. Counter-contributions suggest that the information content and the contribution of a nucleotide are not necessarily correlated. Visualizing learned motifs by aligning the activating sequences (25) would not fully describe how the filter reacts to presented data. It seems that the assumption of nucleotide independence – which is crucial for treating DeepLIFT as a method of estimating Shapley values for input nucleotides (45) – does not hold in full. Indeed, *k*-mer distribution profiles are frequently used features for modelling DNA sequences (as shown also by the dimer-shuffling method of generating reference sequences proposed by Shrikumar et al. (43)). However, DeepLIFT’s multiple successful applications in genomics indicate that the assumption probably holds approximately. We see information content and DeepLIFT’s contribution values as two complementary channels that can be jointly visualized for better interpretability and explainability of CNNs in genomics. Filter enrichment analysis enables even deeper insight in the inner workings of the networks. We generate activation data for hundreds to thousands of species, genes and filters. Yet, aggregation and interpretation of those results beyond case studies is non-trivial, and a promising avenue for further research.

### Genome-scale interpretability

Mapping predictions back to a target genome can be used both as a way of investigating a given model’s performance and as a method of genome analysis. GWPA plots of well-annotated genomes highlight the sequences with erroneous and correct phenotype predictions at both genome and gene level, and nucleotide-resolution contribution maps help track those regions down to individual amino-acids. On the other hand, once a trusted model is developed, it can be used on newly emerging pathogens, as the SARS-CoV-2 virus briefly analyzed in this work. Therefore, we see GPWA applications in both probing the behaviour of artificial neural networks in pathogen genomics and finding regions of interest in weakly annotated genomes. What is more, the approach could be easily co-opted to genome-wide activation analyses of any arbitrary, intermediate neuron. The methods presented here may also be applied to other biological problems, and extending them to other hosts and pathogen groups, multi-class classification or gene identification is possible. However, experimental work and traditional sequence analysis are required to truly understand the biology behind host adaptation and distinguish true hits from false positives.

## Conclusion

We presented a new approach for predicting a host of a novel virus based on a single DNA read or a read pair, cutting the error rates in half compared to the previous state-of-the-art. For convolutional filters, we jointly visualize nucleotide contributions and information content. Finally, we use GWPA plots to gain insights into the models’ behaviour and analyze a recently emerged SARS-CoV-2 virus. The approach presented here is implemented as a python package (see Data availability) and a command line tool easily installable with Bioconda (86).

## DATA AVAILABILITY

The datasets of simulated reads with associated metadata are hosted at https://doi.org/10.5281/zenodo.4312525. The tool can be installed with Bioconda (conda install deepacvir, requires setting up Bioconda), Docker (docker pull dacshpi/deepac) or pip (pip install deepacvir). Detailed installation instructions, user guide and the main codebase (including the interpretability workflows presented here) are available at https://gitlab.com/dacs-hpi/DeePaC. Source code of the plugin shipping the trained models, config files describing the architectures used and the models themselves are available at https://gitlab.com/dacs-hpi/DeePaC-vir.

## ACKNOWLEDGEMENTS

We gratefully acknowledge Yong-Zhen Zhang and the scientists at the Shanghai Public Health Clinical Center & School of Public Health, Fudan University, who shared the sequence of the SARS-CoV-2 virus ahead of publication. We thank Melania Nowicka (Max Plank Institute for Molecular Genetics) for inspiring discussions on efficient calculations of partial Shapley values, Vitor C. Piro (Hasso Plattner Institute) for discussions on traversing taxonomy graphs, Lothar H. Wieler (Robert Koch Institute) for useful comments on the first draft of the manuscript and the Anonymous Reviewers for their suggestions and feedback.

## REFERENCES

1. Calvignac-Spencer, S., Schulze, J. M., Zickmann, F., and Renard, B. Y. (2014) Clock rooting further demonstrates that Guinea 2014 EBOV is a member of the Zaïre lineage. PLoS currents, 6.

2. Vouga, M. and Greub, G. (January, 2016) Emerging bacterial pathogens: the past and beyond. Clinical Microbiology and Infection, 22(1), 12–21.

3. Trappe, K., Marschall, T., and Renard, B. Y. (September, 2016) Detecting horizontal gene transfer by mapping sequencing reads across species boundaries. Bioinformatics, 32(17), i595–i604.

4. Leendertz, S. A. J., Gogarten, J. F., Düx, A., Calvignac-Spencer, S., and Leendertz, F. H. (Mar, 2016) Assessing the Evidence Supporting Fruit Bats as the Primary Reservoirs for Ebola Viruses. EcoHealth, 13(1), 18– 25.

5. Lecuit, M. and Eloit, M. (2014) The diagnosis of infectious diseases by whole genome next generation sequencing: a new era is opening. Frontiers in Cellular and Infection Microbiology, 4, 25.

6. Calistri, A. and Palù, G. (2015) Editorial commentary: Unbiased next-generation sequencing and new pathogen discovery: undeniable advantages and still-existing drawbacks. Clinical Infectious Diseases: An Official Publication of the Infectious Diseases Society of America, 60(6), 889–891.

7. Andrusch, A., Dabrowski, P. W., Klenner, J., Tausch, S. H., Kohl, C., Osman, A. A., Renard, B. Y., and Nitsche, A. (2018) PAIPline: pathogen identification in metagenomic and clinical next generation sequencing samples. Bioinformatics, 34(17), i715–i721.

8. Herfst, S., Schrauwen, E. J. A., Linster, M., Chutinimitkul, S., Wit, E. d., Munster, V. J., Sorrell, E. M., Bestebroer, T. M., Burke, D. F., Smith, D. J., Rimmelzwaan, G. F., Osterhaus, A. D. M. E., and Fouchier, R. A. M. (June, 2012)Airborne Transmission of Influenza A/H5N1 Virus Between Ferrets. Science, 336(6088), 1534–1541.

9. Imai, M., Watanabe, T., Hatta, M., Das, S. C., Ozawa, M., Shinya, K., Zhong, G., Hanson, A., Katsura, H., Watanabe, S., Li, C., Kawakami, E., Yamada, S., Kiso, M., Suzuki, Y., Maher, E. A., Neumann, G., and Kawaoka, Y. (June, 2012) Experimental adaptation of an influenza H5 HA confers respiratory droplet transmission to a reassortant H5 HA/H1N1 virus in ferrets. Nature, 486(7403), 420–428.

10. Lipsitch, M. and Inglesby, T. V. (December, 2014) Moratorium on Research Intended To Create Novel Potential Pandemic Pathogens. mBio, 5(6).

11. Noyce, R. S., Lederman, S., and Evans, D. H. (January, 2018) Construction of an infectious horsepox virus vaccine from chemically synthesized DNA fragments. PLOS ONE, 13(1), e0188453.

12. Thiel, V. (2018) Synthetic viruses-Anything new?. PLoS pathogens, 14(10), e1007019.

13. Edwards, R. A., McNair, K., Faust, K., Raes, J., and Dutilh, B. E. (2016) Computational approaches to predict bacteriophage-host relationships. FEMS microbiology reviews, 40(2), 258–272.

14. Eng, C. L., Tong, J. C., and Tan, T. W. (2014) Predicting host tropism of influenza A virus proteins using random forest. BMC Medical Genomics, 7(3), S1.

15. Xu, B., Tan, Z., Li, K., Jiang, T., and Peng, Y. (July, 2017) Predicting the host of influenza viruses based on the word vector. PeerJ, 5, e3579.

16. Li, H. and Sun, F. (2018) Comparative studies of alignment, alignment-free and SVM based approaches for predicting the hosts of viruses based on viral sequences. Scientific Reports, 8(1), 10032.

17. Mock, F., Viehweger, A., Barth, E., and Marz, M. (08, 2020) VIDHOP, viral host prediction with Deep Learning. Bioinformatics, btaa 705.

18. Gałan, W., Bąk, M., and Jakubowska, M. (2019) Host Taxon Predictor - A Tool for Predicting Taxon of the Host of a Newly Discovered Virus. Scientific Reports, 9(1), 3436.

19. Babayan, S. A., Orton, R. J., and Streicker, D. G. (November, 2018) Predicting reservoir hosts and arthropod vectors from evolutionary signatures in RNA virus genomes. Science, 362(6414), 577–580.

20. Zhang, Z., Cai, Z., Tan, Z., Lu, C., Jiang, T., Zhang, G., and Peng, Y. (2019) Rapid identification of human-infecting viruses. Transboundary and Emerging Diseases, 66(6), 2517–2522.

21. Poplin, R., Chang, P.-C., Alexander, D., Schwartz, S., Colthurst, T., Ku, A., Newburger, D., Dijamco, J., Nguyen, N., Afshar, P. T., Gross, S. S., Dorfman, L., McLean, C. Y., and DePristo, M. A. (2018) A universal SNP and small-indel variant caller using deep neural networks. Nature Biotechnology, 36(10), 983–987.

22. Rizzo, R., Fiannaca, A., La Rosa, M., and Urso, A. (June, 2016) Classification Experiments of DNA Sequences by Using a Deep Neural Network and Chaos Game Representation. In Proceedings of the 17th International Conference on Computer Systems and Technologies 2016 New York, NY, USA: Association for Computing Machinery CompSysTech ‘16 pp. 222–228.

23. Löchel, H. F., Eger, D., Sperlea, T., and Heider, D. (January, 2020) Deep learning on chaos game representation for proteins. Bioinformatics, 36(1), 272–279.

24. Bartoszewicz, J. M., Seidel, A., Rentzsch, R., and Renard, B.Y. (07, 2019) DeePaC: predicting pathogenic potential of novel DNA with reverse-complement neural networks. Bioinformatics, 36(1), 81–89.

25. Alipanahi, B., Delong, A., Weirauch, M. T., and Frey, B. J. (2015) Predicting the sequence specificities of DNA- and RNA-binding proteins by deep learning. Nature Biotechnology, 33(8), 831–838.

26. Zhou, J. and Troyanskaya, O. G. (2015) Predicting effects of noncoding variants with deep learning–based sequence model. Nature Methods, 12(10), 931–934.

27. Zeng, H., Edwards, M. D., Liu, G., and Gifford, D. K. (2016) Convolutional neural network architectures for predicting DNA–protein binding. Bioinformatics, 32(12), i121–i127.

28. Quang, D. and Xie, X. (2016) DanQ: a hybrid convolutional and recurrent deep neural network for quantifying the function of DNA sequences. Nucleic Acids Research, 44(11), e107–e107.

29. Kelley, D. R., Snoek, J., and Rinn, J. L. (2016) Basset: learning the regulatory code of the accessible genome with deep convolutional neural networks. Genome Research, 26(7), 990–999.

30. Greenside, P., Shimko, T., Fordyce, P., and Kundaje, A. (2018) Discovering epistatic feature interactions from neural network models of regulatory DNA sequences. Bioinformatics, 34(17), i629–i637.

31. Nair, S., Kim, D. S., Perricone, J., and Kundaje, A. (July, 2019) Integrating regulatory DNA sequence and gene expression to predict genome-wide chromatin accessibility across cellular contexts. Bioinformatics, 35(14), i108–i116.

32. Avsec, Ž., Weilert, M., Shrikumar, A., Alexandari, A., Krueger, S., Dalal, K., Fropf, R., McAnany, C., Gagneur, J., Kundaje, A., and Zeitlinger, J. (August, 2019) Deep learning at base-resolution reveals motif syntax of the cis-regulatory cod. bioRxiv, p. 737981.

33. Mock, F., Viehweger, A., Barth, E., and Marz, M. (2019) Viral host prediction with Deep Learning. bioRxiv, p. 575571.

34. Ren, J., Song, K., Deng, C., Ahlgren, N. A., Fuhrman, J. A., Li, Y., Xie, X., and Sun, F. (June, 2018) Identifying viruses from metagenomic data by deep learning. arXiv:1806.07810 [q-bio], arXiv: 1806.07810.

35. Tampuu, A., Bzhalava, Z., Dillner, J., and Vicente, R. (September, 2019) ViraMiner: Deep learning on raw DNA sequences for identifying viral genomes in human samples. PLOS ONE, 14(9), e0222271.

36. Eraslan, G., Avsec, Ž., Gagneur, J., and Theis, F. J. (July, 2019) Deep learning: new computational modelling techniques for genomics. Nature Reviews Genetics, 20(7), 389–403.

37. Schneider, T. D. and Stephens, R.M. (October, 1990) Sequence logos: a new way to display consensus sequences. Nucleic Acids Research, 18(20), 6097–6100.

38. Crooks, G. E., Hon, G., Chandonia, J.-M., and Brenner, S.E. (June, 2004) WebLogo: a sequence logo generator. Genome Research, 14(6), 1188– 1190.

39. Lanchantin, J., Singh, R., Lin, Z., and Qi, Y. (2016) Deep Motif: Visualizing Genomic Sequence Classifications. CoRR, abs/1605.01133.

40. Lanchantin, J., Singh, R., Wang, B., and Qi, Y. (2017) Deep motif dashboard: visualizing and understanding genomic sequences using deep neural networks. Pacific Symposium on Biocomputing. Pacific Symposium on Biocomputing, 22, 254–265.

41. Sundararajan, M., Taly, A., and Yan, Q. (2016) Gradients of Counterfactuals. CoRR, abs/1611.02639.

42. Jha, A., Aicher, J. K., Singh, D., and Barash, Y. (2019) Improving interpretability of deep learning models: splicing codes as a case study. bioRxiv,.

43. Shrikumar, A., Greenside, P., and Kundaje, A. (August, 2017) Learning Important Features Through Propagating Activation Differences. In Precup, D. and Teh, Y.W.s, (eds.), Proceedings of the 34th International Conference on Machine Learning, International Convention Centre, Sydney, Australia: PMLR Vol. 70 of Proceedings of Machine Learning Research, pp. 3145–3153.

44. Bach, S., Binder, A., Montavon, G., Klauschen, F., Müller, K.-R., and Samek, W. (July, 2015) On Pixel-Wise Explanations for Non-Linear Classifier Decisions by Layer-Wise Relevance Propagation. PLOS ONE, 10(7), e0130140.

45. Lundberg, S. M. and Lee, S.-I. (2017) A Unified Approach to Interpreting Model Predictions. In Guyon, I., Luxburg, U. V., Bengio, S., Wallach, H., Fergus, R., Vishwanathan, S., and Garnett, R., (eds.), Advances in Neural Information Processing Systems 30, pp. 4765–4774 Curran Associates, Inc.

46. Shrikumar, A., Tian, K., Shcherbina, A., Avsec, Ž., Banerjee, A., Sharmin, M., Nair, S., and Kundaje, A. (March, 2019) TF-MoDISco v0.4.2.2-alpha: Technical Note. arXiv:1811.00416 [cs, q-bio, stat], arXiv:1811.00416.

47. Altschul, S. F., Gish, W., Miller, W., Myers, E. W., and Lipman, D. J. (1990) Basic local alignment search tool. Journal of Molecular Biology, 215(3), 403–410.

48. Camacho, C., Coulouris, G., Avagyan, V., Ma, N., Papadopoulos, J., Bealer, K., and Madden, T. L. (December, 2009) BLAST+: architecture and applications. BMC Bioinformatics, 10(1), 421.

49. Wu, F., Zhao, S., Yu, B., Chen, Y.-M., Wang, W., Hu, Y., Song, Z.- G., Tao, Z.-W., Tian, J.-H., Pei, Y.-Y., Yuan, M.-L., Zhang, Y.-L., Dai, F.-H., Liu, Y., Wang, Q.-M., Zheng, J.-J., Xu, L., Holmes, E. C., and Zhang, Y.-Z. (January, 2020) Complete genome characterisation of a novel coronavirus associated with severe human respiratory disease in Wuhan, China. bioRxiv, p. 2020.01.24.919183.

50. Mihara, T., Nishimura, Y., Shimizu, Y., Nishiyama, H., Yoshikawa, G., Uehara, H., Hingamp, P., Goto, S., and Ogata, H. (2016) Linking Virus Genomes with Host Taxonomy. Viruses, 8(3), 66.

51. King, A. M. Q., Adams, M. J., Carstens, E. B., and Lefkowitz, E. J., (eds.) (2012) Virus Taxonomy: Ninth Report of the International Committee on Taxonomy of Viruses, Academic Press, London; Waltham.

52. Lefkowitz, E. J., Dempsey, D. M., Hendrickson, R. C., Orton, R. J., Siddell, S. G., and Smith, D. B. (January, 2018) Virus taxonomy: the database of the International Committee on Taxonomy of Viruses (ICTV). Nucleic Acids Research, 46(D1), D708–D717.

53. Holtgrewe, M. (2010) Mason – A Read Simulator for Second Generation Sequencing Data. Technical Report FU Berlin,.

54. Deneke, C., Rentzsch, R., and Renard, B. Y. (2017) PaPrBaG: A machine learning approach for the detection of novel pathogens from NGS data. Scientific Reports, 7, 39194.

55. Moustafa, A., Xie, C., Kirkness, E., Biggs, W., Wong, E., Turpaz, Y., Bloom, K., Delwart, E., Nelson, K. E., Venter, J. C., and Telenti, (March, 2017) The blood DNA virome in 8,000 humans. PLOS Pathogens, 13(3), e1006292.

56. Gorbalenya, A. E., Baker, S. C., Baric, R. S., de Groot, R. J., Drosten, C., Gulyaeva, A. A., Haagmans, B. L., Lauber, C., Leontovich, M., Neuman, B. W., Penzar, D., Perlman, S., Poon, L. L. M., Samborskiy, D. V., Sidorov, I. A., Sola, I., Ziebuhr, J., and Coronaviridae Study Group of the International Committee on Taxonomy of Viruses (April, 2020) The species Severe acute respiratory syndrome-related coronavirus : classifying 2019-nCoV and naming it SARS-CoV-2. Nature Microbiology, 5(4), 536–544.

57. Simmonds, P. and Aiewsakun, P. (August, 2018) Virus classification – where do you draw the line?. Archives of Virology, 163(8), 2037–2046.

58. Van Regenmortel, M. H. V. (January, 2018) Chapter One - The Species Problem in Virology. In Kielian, M., Mettenleiter, T. C., and Roossinck, M. J., (eds.),Advances in Virus Research, Vol. 100, pp. 1–18 Academic Press.

59. Li, H. and Durbin, R. (2009) Fast and accurate short read alignment with Burrows–Wheeler transform. Bioinformatics, 25(14), 1754–1760.

60. Langmead, B. and Salzberg, S. L. (2012-03) Fast gapped-read alignment with Bowtie 2. Nature methods, 9(4), 357–359.

61. Wood, D. E. and Salzberg, S. L. (2014) Kraken: ultrafast metagenomic sequence classification using exact alignments. Genome Biology, 15(3), R46.

62. Nix, R. and Kantarciouglu, M. (July, 2012) Incentive Compatible Privacy-Preserving Distributed Classification. IEEE Transactions on Dependable and Secure Computing, 9(4), 451–462 Conference Name: IEEE Transactions on Dependable and Secure Computing.

63. Matejczyk, S. and Michalak, T. (2015) Solving Influence Maximization Problem Using Methods from Cooperative Game Theory., Instytut Podstaw Informatyki PAN, Publication Title: k 20533.

64. Thorvaldsdóttir, H., Robinson, J. T., and Mesirov, J. P. (March, 2013) Integrative Genomics Viewer (IGV): high-performance genomics data visualization and exploration. Briefings in Bioinformatics, 14(2), 178– 192.

65. DeLano, W. L. and others (2002) Pymol: An open-source molecular graphics tool. CCP4 Newsletter on protein crystallography, 40(1), 82–92.

66. Yang, Y.-H., Jiang, Y.-L., Zhang, J., Wang, L., Bai, X.-H., Zhang, S.-J., Ren, Y.-M., Li, N., Zhang, Y.-H., Zhang, Z., Gong, Q., Mei, Y., Xue, T., Zhang, J.-R., Chen, Y., and Zhou, C.-Z. (June, 2014) Structural Insights into SraP-Mediated Staphylococcus aureus Adhesion to Host Cells. PLOS Pathogens, 10(6), e1004169.

67. Stojkova, P., Spidlova, P., and Stulik, J. (2019) Nucleoid-Associated Protein HU: A Lilliputian in Gene Regulation of Bacterial Virulence. Frontiers in Cellular and Infection Microbiology, 9, 159.

68. Li, F. (2016) Structure, Function, and Evolution of Coronavirus Spike Proteins. Annual Review of Virology, 3(1), 237–261.

69. Marchler-Bauer, A., Bo, Y., Han, L., He, J., Lanczycki, C. J., Lu, S., Chitsaz, F., Derbyshire, M. K., Geer, R. C., Gonzales, N. R., Gwadz, M., Hurwitz, D. I., Lu, F., Marchler, G. H., Song, J. S., Thanki, N., Wang, Z., Yamashita, R. A., Zhang, D., Zheng, C., Geer, L. Y., and Bryant, S. H. (2017) CDD/SPARCLE: functional classification of proteins via subfamily domain architectures. Nucleic Acids Research, 45(D1), D200–D203.

70. Wrapp, D., Wang, N., Corbett, K. S., Goldsmith, J. A., Hsieh, C.-L., Abiona, O., Graham, B. S., and McLellan, J. S. (March, 2020) Cryo-EM structure of the 2019-nCoV spike in the prefusion conformation. Science, 367(6483), 1260–1263 Publisher: American Association for the Advancement of Science Section: Report.

71. Yuan, M., Wu, N. C., Zhu, X., Lee, C.-C. D., So, R. T. Y., Lv, H., Mok, C. K. P., and Wilson, I. A. (May, 2020) A highly conserved cryptic epitope in the receptor binding domains of SARS-CoV-2 and SARS-CoV. Science, 368(6491), 630–633 Publisher: American Association for the Advancement of Science Section: Report.

72. Starr, T. N., Greaney, A. J., Hilton, S. K., Crawford, K. H., Navarro, M. J., Bowen, J. E., Tortorici, M. A., Walls, A. C., Veesler, D., and Bloom, J. D. (June, 2020) Deep mutational scanning of SARS-CoV-2 receptor binding domain reveals constraints on folding and ACE2 binding. bioRxiv, p. 2020.06.17.157982 Publisher: Cold Spring Harbor Laboratory Section: New Results.

73. Pinto, D., Park, Y.-J., Beltramello, M., Walls, A. C., Tortorici Jaconi, M. A., Bianchi, S., S., Culap, K., Zatta, F., De Marco, A., Peter, A., Guarino, B., Spreafico, R., Cameroni, E., Case, J.B., Chen, R.E., Havenar-Daughton, C., Snell, G., Telenti, A., Virgin, H. W., Lanzavecchia, A., Diamond, M.S., Fink, K., Veesler D.s, and Corti, D. (May, 2020) Cross-neutralization of SARS-CoV-2 by a human monoclonal SARS-CoV antibody. Nature, pp. 1–10 Publisher: Nature Publishing Group.

74. Brendel, W. and Bethge, M. (2019) Approximating CNNs with Bag-of-local-Features models works surprisingly well on ImageNet. In International Conference on Learning Representations.

75. National Research Council (2010) Sequence-Based Classification of Select Agents: A Brighter Line, The National Academies Press, .

76. National Academies of Sciences, Engineering, and Medicine (2018) Biodefense in the Age of Synthetic Biology, The National Academies Press,

77. Diggans, J. and Leproust, E. (2019) Next Steps for Access to Safe, Secure DNA Synthesis. Frontiers in Bioengineering and Biotechnology, 7.

78. O’Brien, J. T. and Nelson, C. (June, 2020) Assessing the Risks Posed by the Convergence of Artificial Intelligence and Biotechnology. Health Security, 18(3), 219–227.

79. Brookes, D., Park, H., and Listgarten, J. (May, 2019) Conditioning by adaptive sampling for robust design. In International Conference on Machine Learning pp. 773–782.

80. Alley, E. C., Khimulya, G., Biswas, S., AlQuraishi, M., and Church, G. M. (December, 2019) Unified rational protein engineering with sequence-based deep representation learning. Nature Methods, 16(12), 1315–1322.

81. Biswas, S., Khimulya, G., Alley, E. C., Esvelt, K. M., and Church, G. M. (January, 2020) Low-N protein engineering with data-efficient deep learning. bioRxiv, p. 2020.01.23.917682.

82. Gupta, A. and Zou, J. (February, 2019) Feedback GAN for DNA optimizes protein functions. Nature Machine Intelligence, 1(2), 105–111.

83. Gupta, A. and Kundaje, A. (July, 2019) Targeted optimization of regulatory DNA sequences with neural editing architectures. bioRxiv, p. 714402.

84. Linder, J., Bogard, N., Rosenberg, A. B., and Seelig, G. (December, 2019) Deep exploration networks for rapid engineering of functional DNA sequences. bioRxiv, p. 864363.

85. Schreiber, J., Lu, Y. Y., and Noble, W. S. (May, 2020) Ledidi: Designing genomic edits that induce functional activity. bioRxiv, p. 2020.05.21.109686.

86. Grüning, B., Dale, R., Sjödin, A., Chapman, B. A., Rowe, J., Tomkins-Tinch, C. H., Valieris, R., and Köster, J. (July, 2018) Bioconda: sustainable and comprehensive software distribution for the life sciences. Nature Methods, 15(7), 475–476 Number: 7 Publisher: Nature Publishing Group.

