## Supplementary Information for "Interpretable detection of novel human viruses from genome sequencing data"

Jakub M. Bartoszewicz[1 2 3 4], Anja Seidel[1 2 5] and Bernhard Y. Renard[1 3 4]

Table S1: Classification accuracy depending on the negative class definition, read pairs. Euk. – Eukaryota dataset; Met. – Metazoa dataset, Cho. – Chordata dataset, Str. – Stratified dataset, X-150 – first 150 bp of each read in X. Training set in subscript of the model name; test set in column headers. Recall (Rec.) is identical in all cases, as the positive class remains unchanged. Best performance in bold. $CNN_{All}$ achieves best overall accuracy.

| | All | Euk. | Met. | Cho. | Str. | Rec. |
|---|---|---|---|---|---|---|
| $CNN_{ALL}$ | **89.9** | 85.9 | 83.3 | 78.6 | **88.1** | **85.4** |
| $CNN_{CHO}$ | 84.9 | 84.2 | 83.6 | 82.4 | 84.6 | 71.7 |
| $CNN_{STR}$ | 88.2 | **86.4** | **85.1** | **82.7** | 87.4 | 78.8 |
| $CNN_{ALL\text{-}150}$ | 89.4 | 85.5 | 82.9 | 78.4 | 87.7 | 83.2 |
| $CNN_{STR\text{-}150}$ | 88.2 | 86.3 | 84.9 | 82.5 | 87.3 | 78.3 |
| $LSTM_{ALL}$ | 86.4 | 78.2 | 74.1 | 65.5 | 82.6 | 83.0 |
| $LSTM_{CHO}$ | 82.8 | 81.9 | 80.8 | 80.0 | 82.4 | 70.6 |
| $LSTM_{STR}$ | 85.8 | 82.2 | 79.6 | 75.2 | 84.2 | 76.3 |

[1]Bioinformatics (MF1), Department of Methodology and Research Infrastructure, Robert Koch Institute, Berlin, Germany
[2]Department of Mathematics and Computer Science, Free University of Berlin, Berlin, Germany
[3]Hasso Plattner Institute for Digital Engineering, Potsdam, Brandenburg, Germany
[4]Digital Engineering Faculty, University of Postdam, Potsdam, Brandenburg, Germany
[5]Currently at: Central Research Institute of Ambulatory Health Care, Berlin, Germany

Table S2: Classification performance in the fully open-view setting (all virus hosts), single reads. Acc. – accuracy, Prec. – precision, Rec. – recall, Spec. – specificity. BLAST yields no predictions for over 12% of the samples. Best performance in bold.

| | Acc. | Prec. | Rec. | Spec. |
|---|---|---|---|---|
| $\text{CNN}_{\text{All}}$ (ours) | **87.8** | 89.9 | **85.2** | **90.4** |
| $\text{LSTM}_{\text{All}}$ (ours) | 84.7 | 86.0 | 82.8 | 86.5 |
| $k$-NN | 75.5 | 76.3 | 73.9 | 77.1 |
| BLAST | 78.4 | **98.3** | 79.2 | 77.6 |

Table S3: Classification performance, novel species of Chordata-infecting viruses. Top: paired reads. BLAST yields predictions for only 56.6% of the pairs. Bottom: whole available genomes or contigs – negative class is the majority class. BAcc. – balanced accuracy (equal to accuracy for the balanced paired-read dataset), Rec. – recall, Spec. – specificity. BLAST (reads) and our networks use read-wise majority vote or output averaging to aggregate predictions over all reads from a genome. BLAST (genome) uses contig-wise majority vote. BLAST (contigs) represents performance on individual contigs treated as separate entities. Note that low precision is heavily affected by class imbalance.

| | BAcc. | Prec. | Rec. | Spec. |
|---|---|---|---|---|
| $\text{CNN}_{\text{SP-Cho}}$ (ours) | **58.2** | 60.6 | 47.1 | **69.3** |
| $\text{CNN}_{\text{SP-All}}$ (ours) | 57.4 | 56.5 | **65.0** | 49.9 |
| BLAST | 37.1 | **65.8** | 19.1 | 55.1 |
| $\text{CNN}_{\text{SP-Cho}}$ (ours) | 54.0 | 42.3 | 22.9 | **85.0** |
| $\text{CNN}_{\text{SP-All}}$ (ours) | 57.4 | 37.9 | **69.8** | 45.0 |
| BLAST (reads) | 57.5 | **47.9** | 35.4 | 79.5 |
| BLAST (genome) | **58.0** | 49.3 | 38.5 | 77.5 |
| BLAST (contigs) | 51.6 | 42.2 | 37.1 | 66.2 |

Table S4: Gene ranking for *S. aureus* (top 3 out of 870). hupB is indirectly engaged in virulence. Our method detects functionally relevant genes using the original DeePaC RC-CNN model.

| Rank | Gene | Score | Biological Process |
|---|---|---|---|
| 1 | sarR | 0.644 | **Virulence** |
| 2 | hupB | 0.642 | DNA condensation |
| 3 | sspB | 0.637 | **Virulence** |

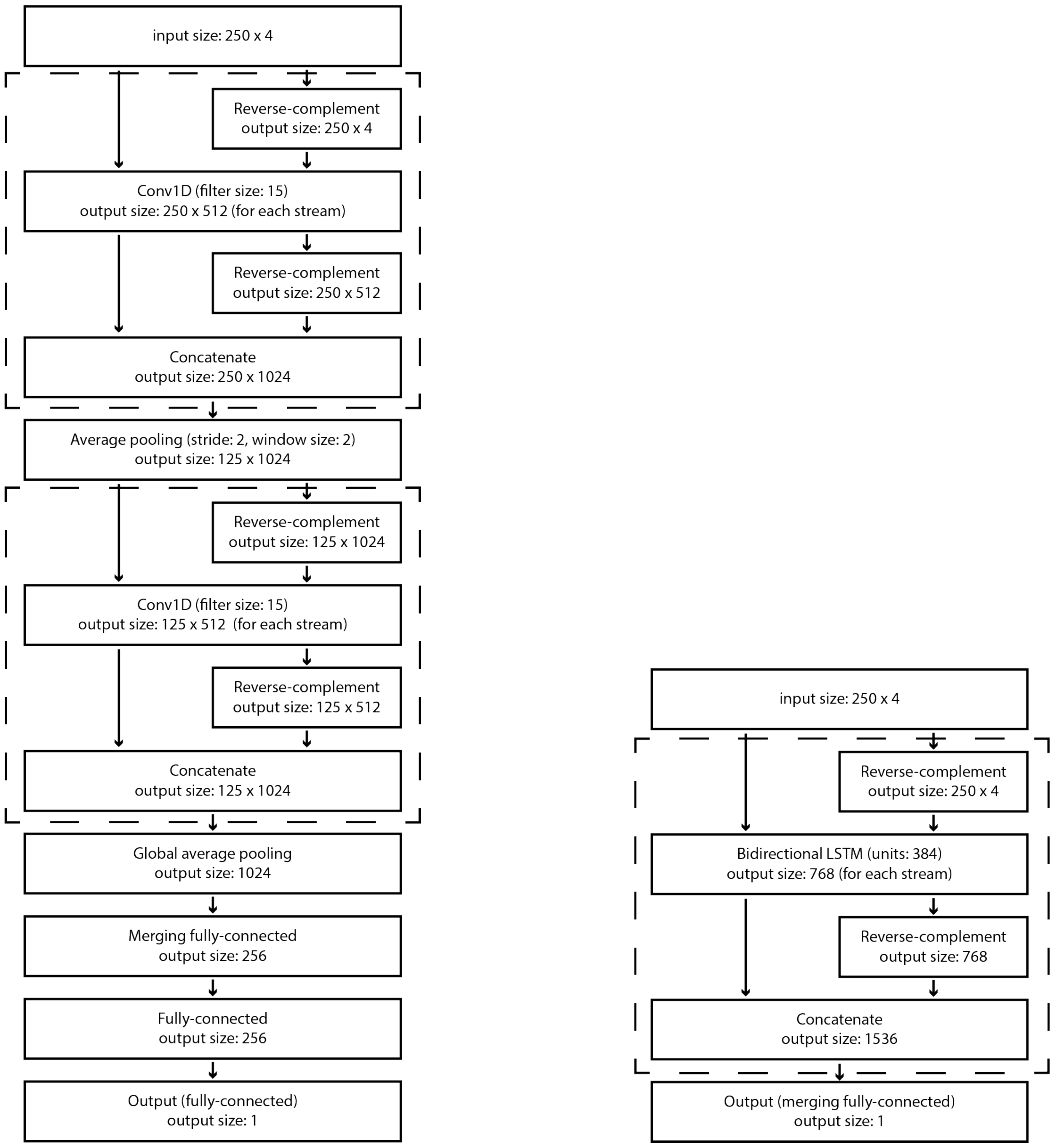


Figure S1: Simplified plots of the *full* reverse-complement architectures used in this study. We omit the batch dimension for clarity. We use sigmoid activation for the output layer and ReLU activation after fully-connected (FC) layers and RC-Convolution blocks (S1a, dashed lines). The RC-LSTM blocks (S1b, dashed lines) use the default combination of activation functions for the LSTM layers as implemented in the `tf.keras` library. We also use dropout after the input (p=0.25), convolutional, recurrent and FC layers (p=0.5). Padding is handled automatically by Keras so that the input length does not change after convolutions. Note that the nucleotide dimension is the channel dimension (the input shape is sequence length x number of channels), and we use 1D convolutions. Hence, convolutions, padding and pooling have no vertical dimension. The merging FC layers sum representations for corresponding channels in the "forward" and "reverse-complement" orientation. Virus-level models were trained on Tesla P100 GPUs using early stopping with a patience of 10 epochs for a default maximum of 14 epochs (corresponding to maximum 80h wall-time on this GPU). Species-level model were trained on Tesla V100 GPUs; as convergence took longer we allowed a maximum of 160h (corresponding to a triple increase in maximum epochs). A trained model can be used for either single reads or read pairs; in the latter case we run predictions separately for both mates and the average the final outputs. More in-depth description of the reverse-complement architectures can be found in Bartoszewicz *et al.* (2019). The architectures presented in Fig. S1a and Fig. S1b can be reproduced using the `deepac-vir train -r` and `deepac-vir train -s` commands, respectively; input data can be supplied using the `-T`, `-t`, `-V` and `-v` flags. Config files can be retrieved using `deepac-vir templates`, modified and used with `deepac train -c` to train custom models. See user guide for details.

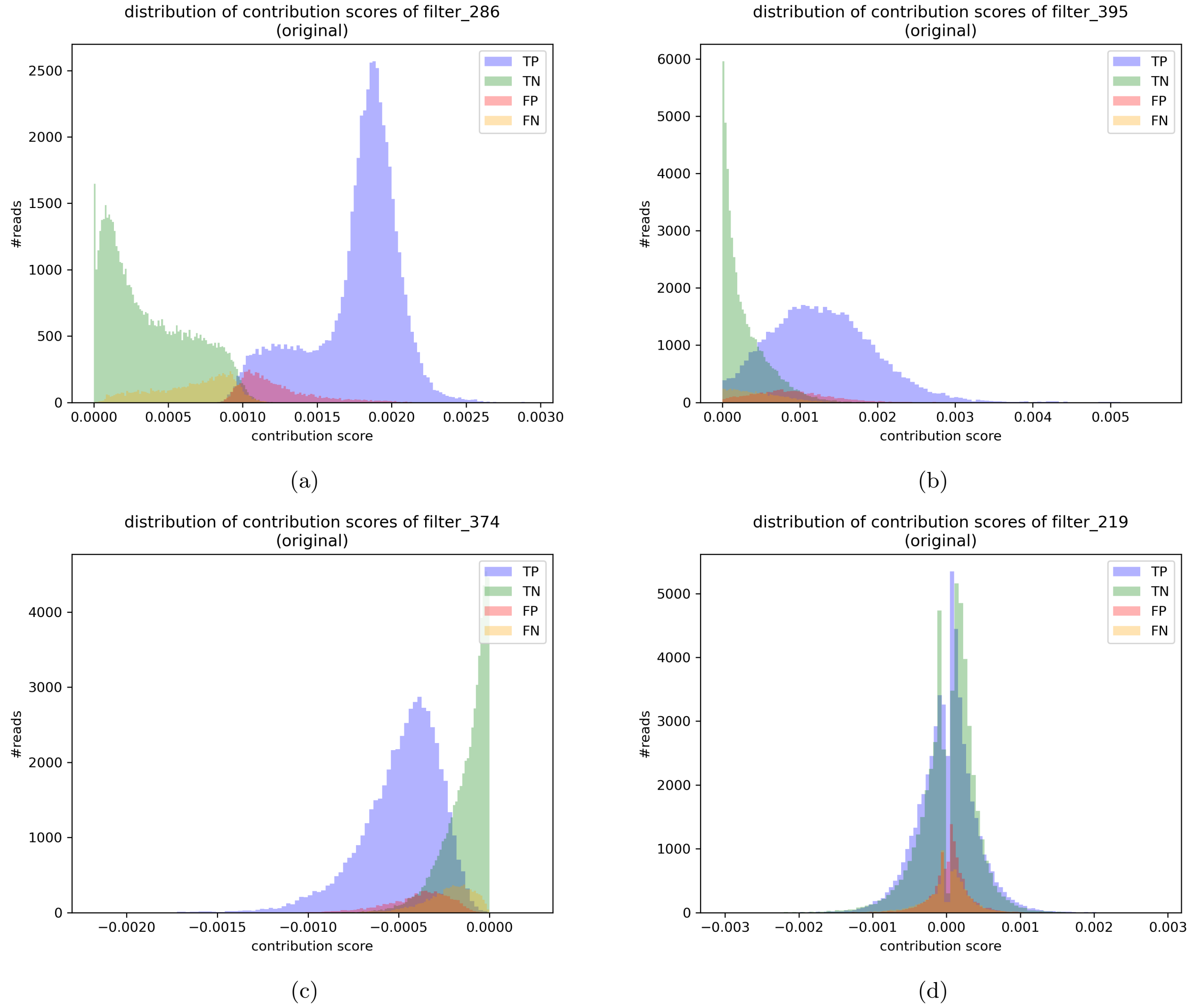


Figure S2: Contribution histograms of selected filters. S2a: The ”ambiguity detector” with the highest mean contribution score. S2b-S2d: Filters presented in Fig. 1 (main text). S2b: Second-highest mean contribution score. Contributions are higher for the positive class. S2c: Lowest mean contribution score. Positive class reads are ”penalized” with stronger negative contributions than the negative class reads. S2d: Oscillating filter from the original, bacterial DeePaC RC-CNN. Symmetric contribution distribution suggests that the contributions of the filter are context-dependent, although positive on average.

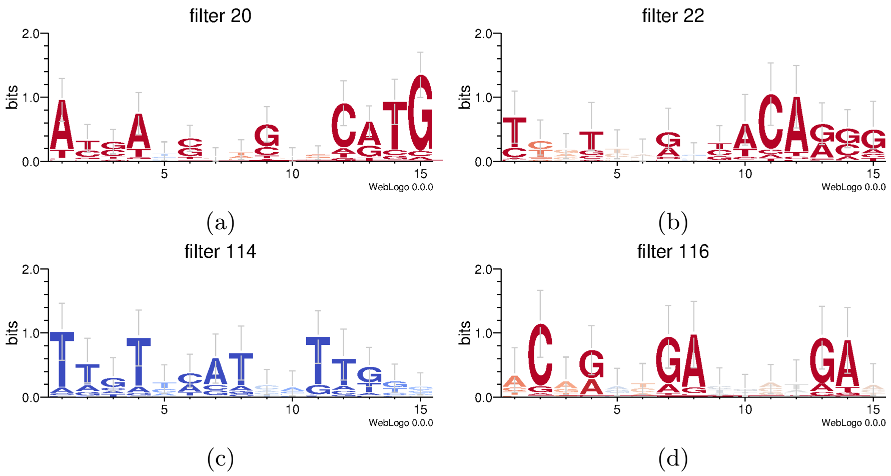


Figure S3: Example oscillating filters with a codon-like structure, extracted from the $\text{CNN}_{\text{All}}$ model. Consensus sequences: AYSANSNNGNNCATG (S3a), TYNTNNNRNNACARSG (S3b), TTGTNMATNNNTTKNN (S3c), ACNRNNGANNNNNGAN (S3d).

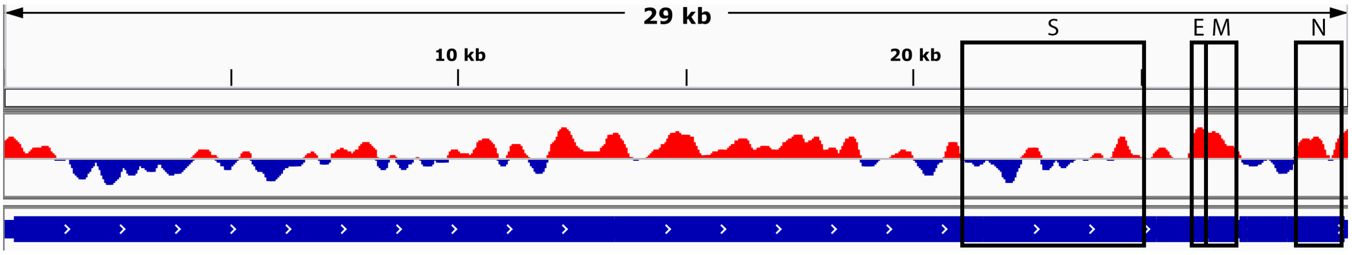


Figure S4: GWPA plot of the SARSr-CoV RaTG13 virus scored by the $\text{CNN}_{\text{All}}$ model. The pattern of predicted infectious potentials is roughly similar to its relative, SARS-CoV-2. Regions of elevated infectious on the right end of the plot correspond to E, M, and N genes; the wide red region in the middle is collocated with ORF1b. Homologous regions in SARS-CoV-2 are scored highly as well.

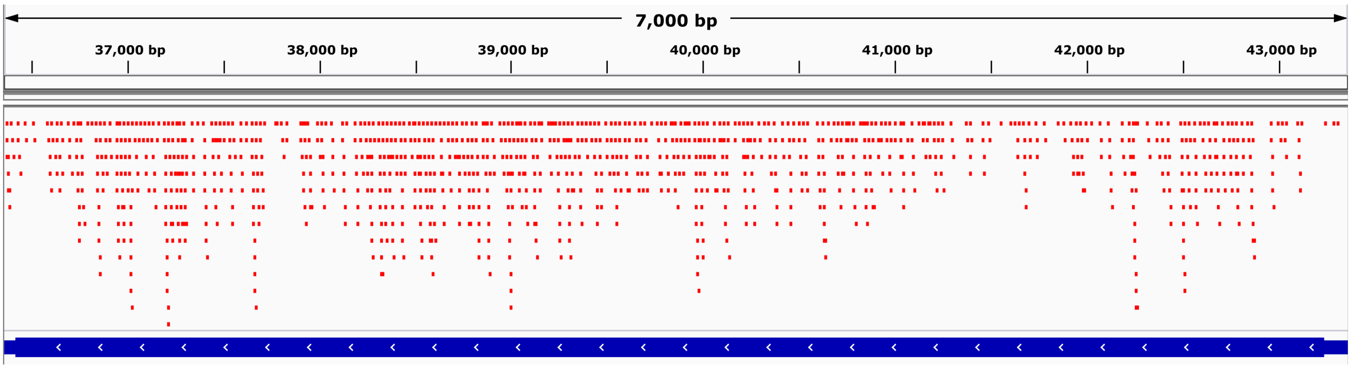


(a)

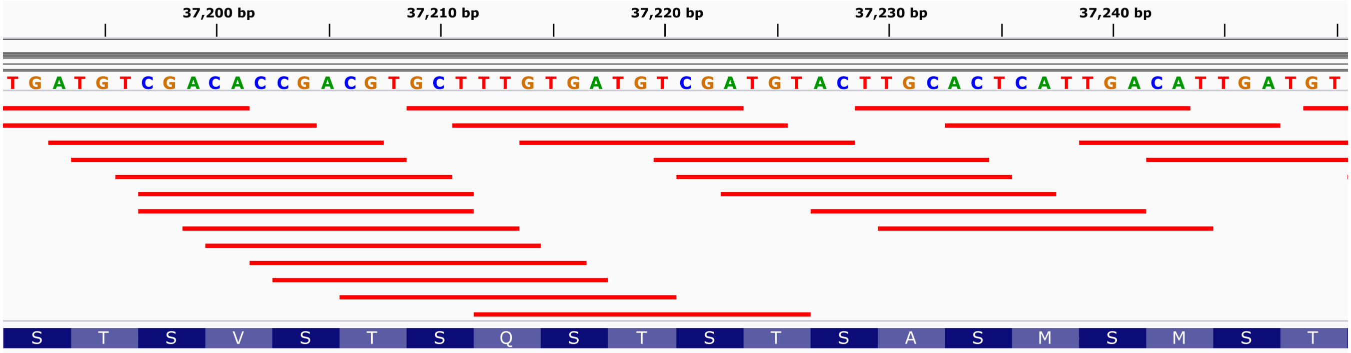


(b)

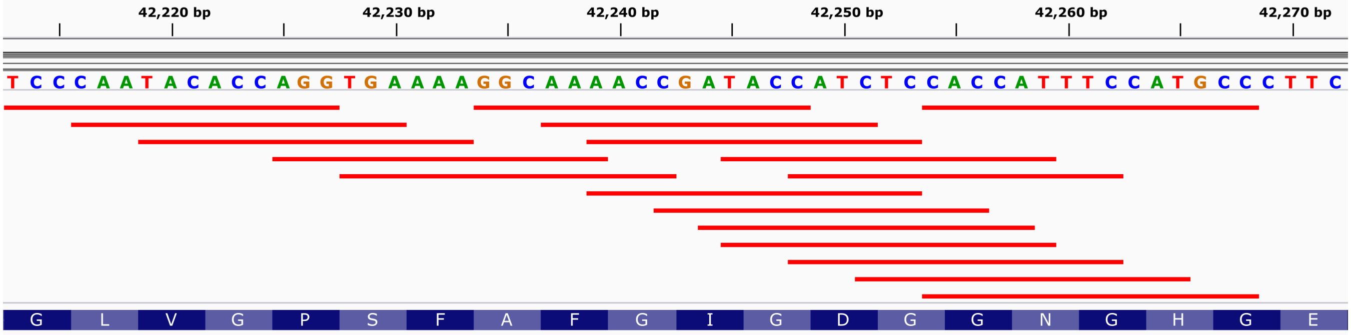


(c)

Figure S5: Activations of filter 219 (original DeePaC RC-CNN) in the sraP gene of *S. aureus* subsp. *aureus* 21200. Top, in red: filter hits. Bottom, in blue: reference sequence annotation (contig 19). S5a: sraP, a serine-rich adhesin for platelets, the top hit gene in the enrichment analysis for filter 219. Serine-rich repeat regions cover over 71% of the sequence. S5b: An example stretch of filter 219 activations in a serine-rich repeat region. The filter seems to detect the serine repeats. S5c: An example stretch of filter 219 activations in a non-repeat region. The filter seems to detect a local glycine repeat.

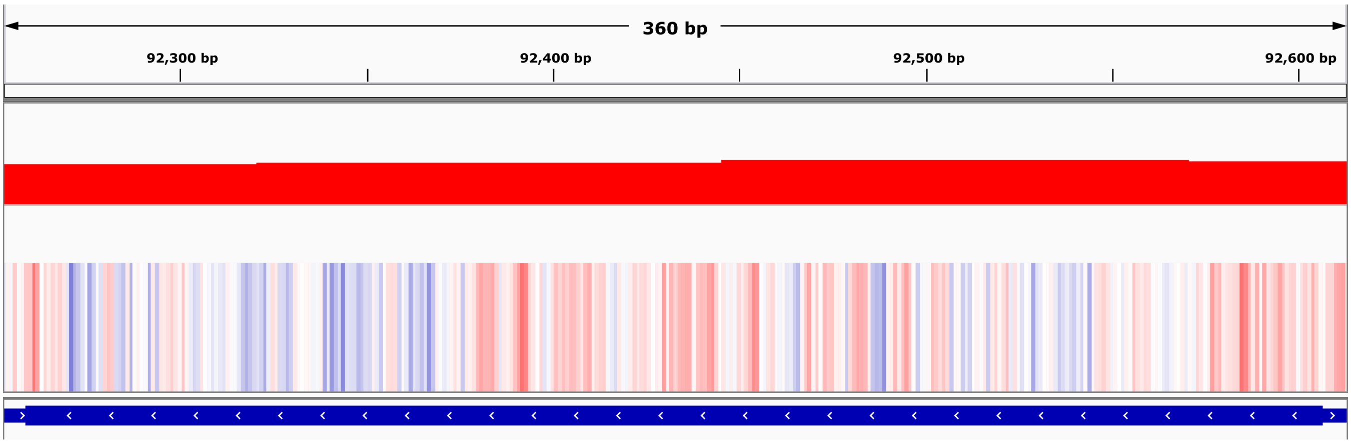


(a)

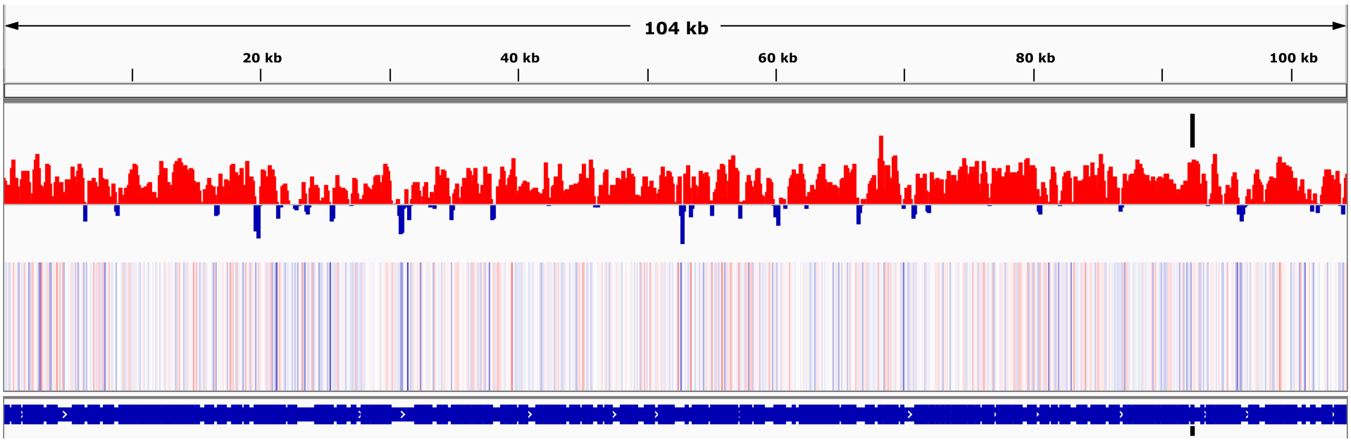


(b)

Figure S6: GWPA of *S. aureus* subsp. *aureus* 21200. Top: score predicted by the original DeePaC RC-CNN. Heatmap: nucleotide contributions. Bottom, in blue: reference sequence annotation. Negative contributions marked with blue, positive contributions in red. S6a: sarR gene, with the highest average pathogenicity score. S6b: Contig 18, with sarR location marked with black bars. Neighbouring higher peaks belong to putative genes lacking ground truth annotation.

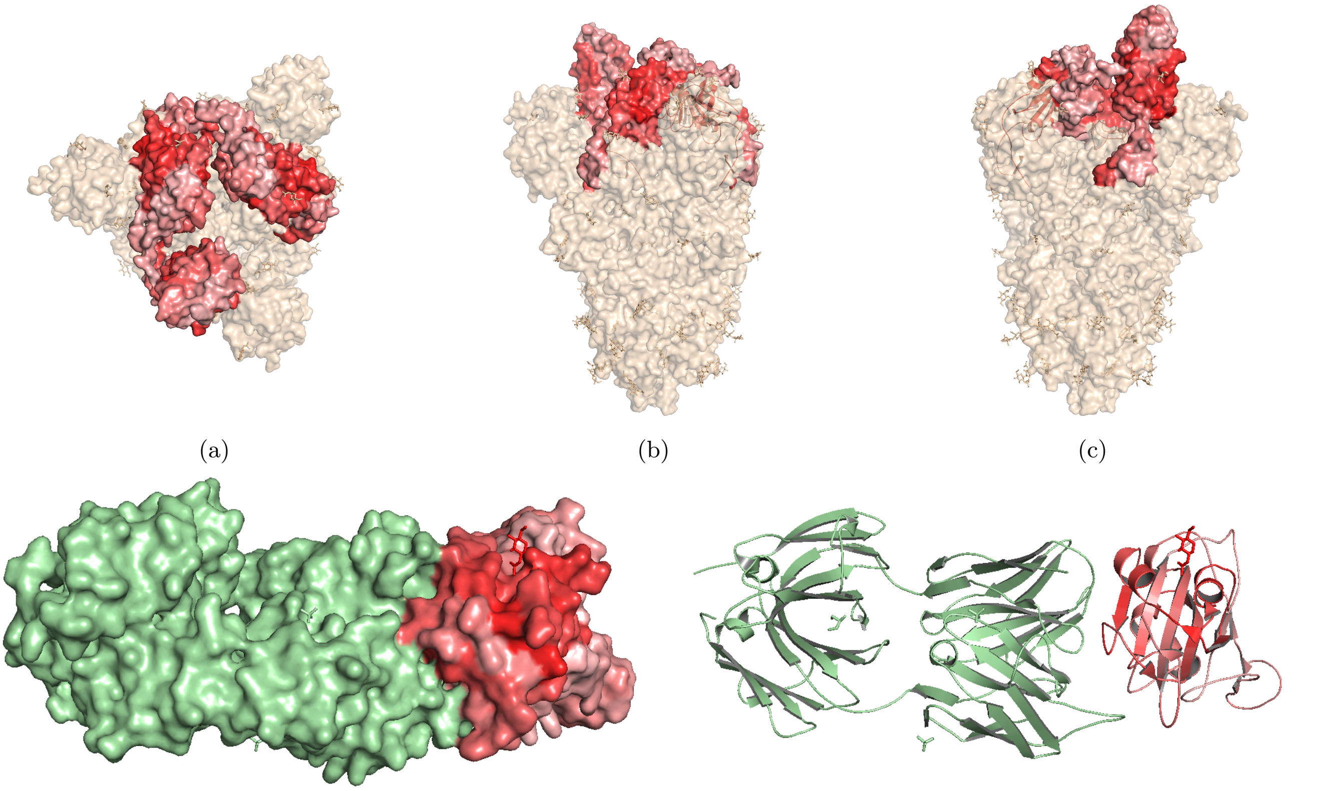

(a) (b) (c) (d) (e)

Figure S7: This figure is analogous to Fig. 3 (main text), but presents average nucleotide contribution scores (per residue) instead of the average predicted pathogenic potential. S7a-S7c: Top and side view of the SARS-CoV-2 spike glycoprotein. Three receptor-binding domains (RBDs) are colored according to the per-residue average contribution score of the corresponding genomic sequence. Since all contributions are positive, the color gradient is dominated by different shades of red. One of the domains is in the "up" conformation. Darker red regions corresponding to the peak in Fig. 2c (main text) are located in the core-RBD subdomain. S7d: RBD in complex with a SARS-neutralizing antibody CR3022 (green). The darker red region covers over 70% of the CR3022 epitope, but spans also to the neighboring fragments, including the N343 glucosaminylation site (carbohydrate in red stick representation). This is a part of the epitope of another neutralizing antibody, S309. S7e: Cartoon representation of Fig. S7d. The red region is centered on two exposed α-helices surrounding the core β-sheet (lower score, light red).

### References

Bartoszewicz, J. M. *et al.* (2019). DeePaC: predicting pathogenic potential of novel DNA with reverse-complement neural networks. *Bioinformatics*, **36**(1), 81–89.
